# Functional rescue of critical-size bone defect using molecular network analysis of axolotl limb regeneration

**DOI:** 10.64898/2025.12.10.692203

**Authors:** A. Polikarpova, I. Rivero-Garcia, T. Gerber, J. Wang, M. Novatchkova, A. Fischer, M. Torres, F. Sánchez-Cabo, E.M. Tanaka

## Abstract

The salamander limb has served as a canonical model for successful regeneration that has yielded numerous molecular insights on its basic mechanism. Harnessing such information to induce regeneration in a non-regenerative setting has been a long-sought goal. The amputated salamander limb efficiently regenerates all of its missing bones, but paradoxically, a large bone gap without amputation – commonly called a Critical Size Defect (CSD) – is not regenerated, similarly to other vertebrates^1^. This non-regenerating injury provides a human-relevant setting to understand how to rescue lack-of-regeneration. Satoh and colleagues demonstrated in axolotl that transplantation of blastema cells from an amputated limb into a CSD yields cartilage bridging of the CSD^2^. This work provided a roadmap for rescuing the CSD. Here we asked, what are the crucial molecular differences in cells populating a regenerating blastema versus the CSD and can they be used to elicit CSD bridging? Previous genetic fate mapping and single cell transcriptomics showed that limb regeneration occurs via migration and dedifferentiation of fibroblastic, soft connective tissue (CT) cells that form a multipotent, skeletal stem cell ^3, 4^. Here using single cell transcriptomics and genetic fate mapping we found that the CSD is populated by CT cells that undergo a divergent molecular transition compared to the blastema. Using gene regulatory network (GRN) modeling, and lipid nanoparticle (LNP) delivery of mRNA we could express a single molecular factor, *Wnt3a*, to induce cartilage bridging of the CSD. Our results demonstrate the power of using molecular information from successful regeneration to rescue a non-regenerating injury.

## Main

To compare healing after hindlimb amputation to a critical-sized defect healing, we generated a CSD in the hindlimb by surgically generating a ~3 mm bone gap in the femur that was stabilized using a polyolefin sleeve. Movat pentachrome staining of sectioned tissue showed that over six-weeks, the amputated sample regenerated the tibia, fibula and foot skeleton, whereas in the CSD, no evidence of skeleton formation was observed in the middle of the gap (Figure 1A). To compare the cells populating the CSD versus the blastema we performed a scRNA-seq time course of cells at the injury site in the two models. Cell-type annotation revealed that both samples contained CT cells as well as immune cells, endothelial cells and epidermis (Figure 1B-D, S1). The CSD sample additionally harbored cells annotated as myogenic cells, consistent with histological data in which muscle tissue could be observed in the outer radius of the CSD limb (Figure 1A asterisk, S1C). Genetic-fate-mapping using the double transgenic *Prrx1:TFPnls-T2A-Cre-ERT2;CAGGs:lp-STOP-lp-Cherry* ^4, 5^ (short *Prrx1-LP-Cherry*) revealed that the majority of CT-derived cells in the gap originated from the soft connective tissue with limited contribution from the periskeleton, similar to the regenerating limb blastema^4^ (Figure 1E-G). The lower percentage of CT cells in the CSD at 3 weeks post-injury (wpi) compared to the blastema (Figure 1F) was consistent with the presence of muscle-lineage cells in the CSD as seen in the histological and scRNA-seq datasets. From this data, we conclude that the mesenchymal tissue of the limb blastema and the CSD harbor cells of similar CT origin.

**Figure 1.**
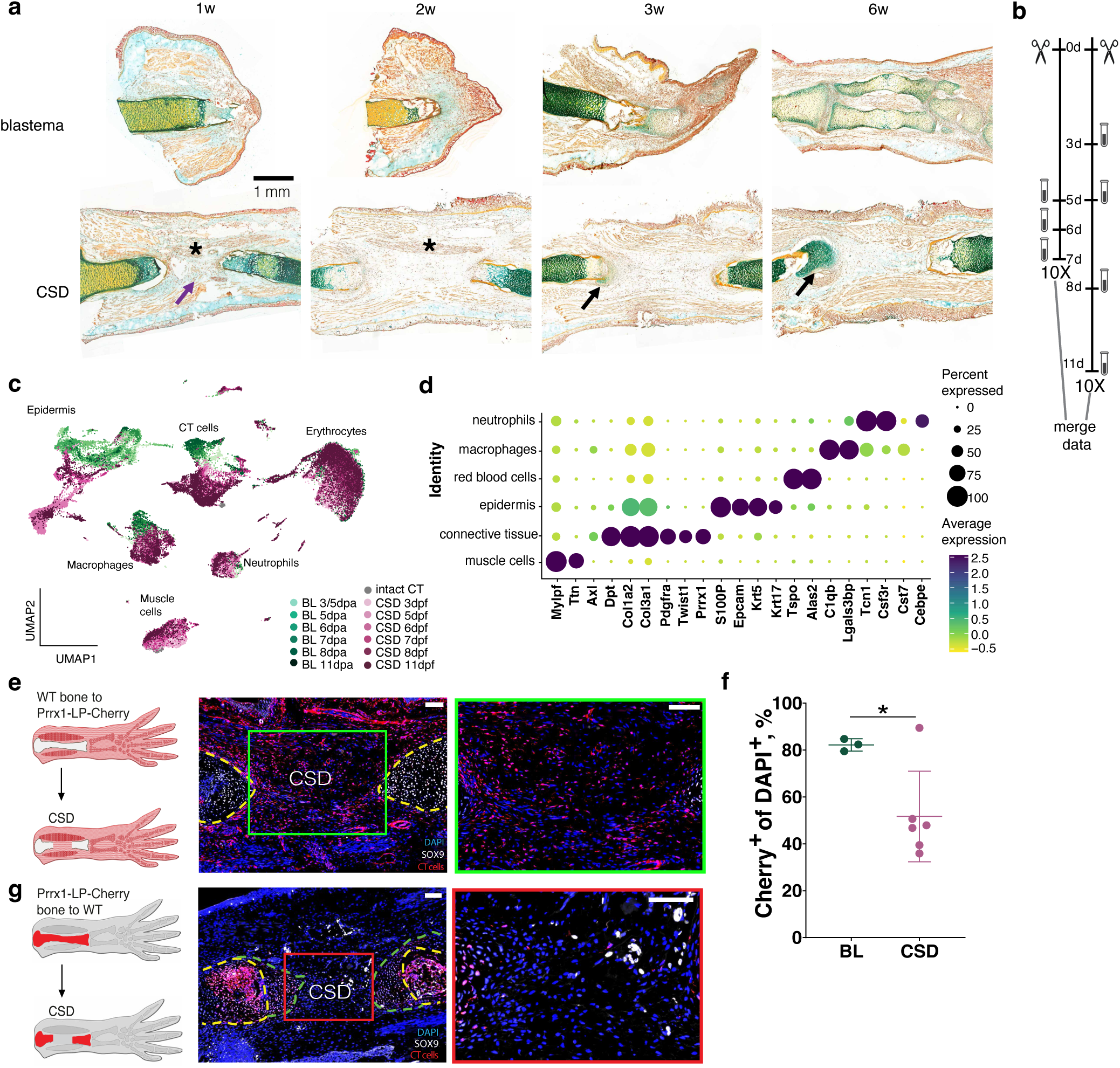
Despite cellular composition similar to blastema, CSD cannot regenerate bone. **a.** Movat pentachrome staining of blastema and critical size femoral fracture in axolotl upper hind limb. The cartilage is stained in blue/green, the ossified bone – in yellow, and the muscle – in red/brown and marked with an asterisk. Purple arrow points at the CSD gap, and black arrows – at the cartilage caps at the CSD edges. Scale bar 1 mm. N = 3. **b.** Schema of the scRNA-seq experiments. Blastema and CSD cells were harvested on 0, 3, 8, and 11 dpi for the 1^st^ batch and on days 5, 6, 7 for the 2^nd^ batch. Data from 2 batches was integrated for the subsequent data analysis, using Harmony^35^. **c.** Uniform manifold approximation and projection (UMAP) atlas of cell classes involved in axolotl limb and bone regeneration based on the 10X Genomics scRNA-seq data. **d.** Dot plot showing gene expression intensity and frequency in the cell clusters from C. **e.** Representative images of wildtype femur-transplanted *Prrx1-LP-Cherry* limbs with CSD. Cartilage cells and precursors express SOX9, and CT progenitors are stained with a-PRRX1 antibody. The right framed panel is a zoom of the left image. Scale bars 200 µm. Bones are marked with yellow dashed lines. **f.** Cherry^+^ cells from soft CT tissue contribute to CSD gap, albeit to a lower extent than to blastema. Data is normalized to the reporter conversion rate in recipient *Prrx1-LP-Cherry* animals. N = 3 for blastema, N = 6 for CSD, with several sections quantified and averaged for each replicate. * p < 0.05. **g.** Representative images of *Prrx1-LP-Cherry* upper hind limb femur-transplanted limbs with CSD. Bone-derived cells were observed in the CSD gap. The right framed panel is a zoom of the left image. Scale bars 200 µm. Calluses are marked with green dashed lines, bones – with yellow dashed lines.

Since previous genetic fate mapping of limb regeneration showed that soft CT-derived cells regenerate patterned skeleton ^3, 4^, we focused our remaining analysis on annotating the CT-derived cells from blastema versus the CSD to understand their underlying differences. Zooming into the CT clusters from both injury datasets by subclustering allowed us to discern emergent cell subtypes associated with the time course of both injuries (Figure 2A, B). In line with the presence of muscle lineage cells in the CSD samples we identified a CSD-specific *Klf5*^+^ CT population that we could associate with muscle fascia (Figure 2A, B). We did not see signs of participation in the regeneration process and thus removed them from the following analyses. In addition, we could identify early blastema and CSD cells at 3 and 5 days post injury (dpi) followed by progenitor cell signatures emerging at 6-8 dpi and then the appearance of cell specification markers for cartilage and tendon at 8-11 dpi (Figure 2A, B). The pseudotime calculation suggested that cell identities between the injuries diverged over time (Figure S2). To assess relative transcriptional similarity between cognate timepoints we pairwise compared the 6 subclusters of CT cells by time point and injury type (Figure 2C, D) and found the lowest correlation between BL and CSD progenitor cell stage samples (6-8 dpi) compared to early cells (3-5 days), or between the late, cartilage clusters derived from both injuries. Previous work showed that regenerative blastema cells acquire the limb bud progenitor identity state after day 5 ^4^, which, in our dataset corresponded to the timepoints (6-8 dpi) – the most divergent transcriptional signatures between blastema and CSD (Figure 2C, D). Gene expression enriched in the blastema progenitors included genes associated with the limb development network such as *Fgf8*, *Grem1, Msx2* along with *Il11* and *Kazald1*, that had previously been associated with the blastema cell state ^4, 6, 7, 8^ (Figure 2E, F, S3A). Notably, *Kazald1* and *Tnc* were expressed across BL cells, while in CSD these genes were enriched in tendon/cartilage population adjacent to the bone edges (Figure 2E, F, S3B, C, D) Conversely, the CSD mesenchyme was enriched for expression of *Fgf2* (Figure 2E, S3A). These data indicated that CT cells that enter the CSD never attain the full limb bud progenitor cell state, and do not activate the FGF8-Shh feedback loop that is required for sustained cell proliferation of blastema cells ^9^.

**Figure 2.**
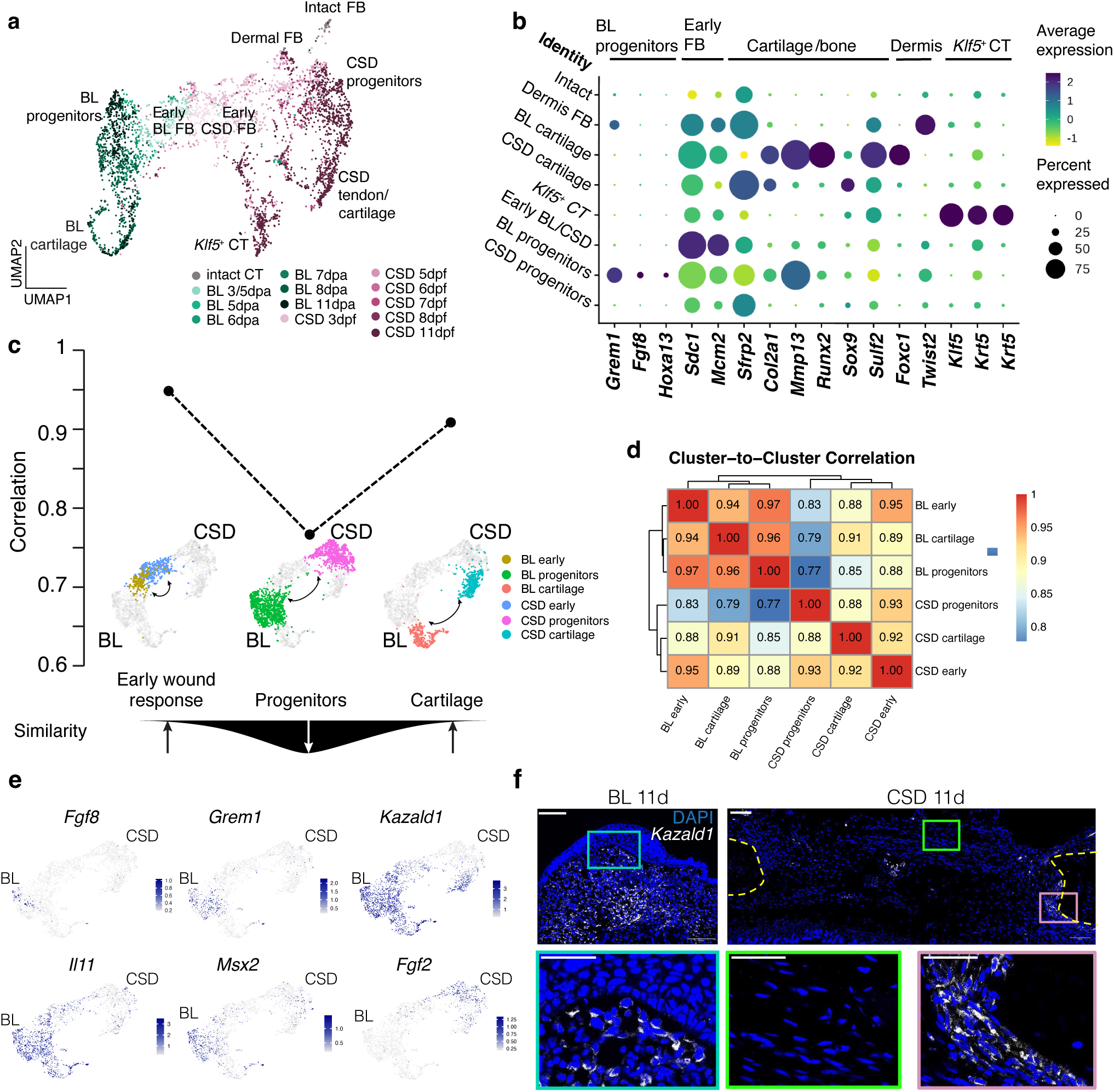
CSD connective tissue cells cannot activate progenitor phenotype. **a.** UMAP embedding of CT cells extracted from the blastema and CSD scRNA-seq data. Blastema cells in green, CSD cells in purple. The color shade indicates harvesting time point. **b.** Dot plot showing gene expression intensity and frequency in the CT cell clusters from A. **c.** Top: Stagewise correlation plot shows lower correlation of intermediate cells from CSD and blastema based on pseudo bulk versus pseudo bulk correlation. Bottom: Summary schematic of the pseudo-bulk correlation result. **d.** Cluster-to-cluster correlation in blastema and CSD CT cells. **e.** Blastema marker expression in CT derived from blastema and CSD. Note prevalence of expression in blastema cells, except for the early blastema marker, *Fgf2*, which is almost exclusively expressed in the CSD-derived CT cells. **f.** HCR *in situ* hybridization was used to visualize and localize the *Kazald1* transcripts (white dots) in blastema and CSD 11 days post-injury. Note absence of *Kazald1* expression in the mid-CSD, and high expression in blastema and the CSD edges. Scale bars: upper row - 200 μm, lower row - 100 μm.

To compare proliferation dynamics, we analyzed the scRNA-seq dataset for cell cycle markers (Figure 3A, S4A). Both early blastema and early CSD cells displayed S-phase-associated transcriptional states. Later, strong S and G2 transcriptional states were found in the blastema progenitor cell states but less in the CSD progenitor states. To directly examine cellular proliferation, we performed pulse labeling with EdU in parallel samples at weekly intervals in *Prrx1-LP-Cherry* transgenic animals (Figure 3B, C). In blastema samples, we observed a peak of EdU incorporation at two wpi with 75% of Cherry^+^ cells incorporating EdU, and 50% of Cherry^+^ cells labeling at three wpi. In contrast, early (one wpi) CSD samples displayed significant EdU incorporation but two and three wpi CSD samples showed very little incorporation (Figure 3C). The results of EdU incorporation assay are mirrored by PCNA staining (Figure S4B). Taken together, the scRNA-seq and EdU labeling studies concur that blastema cells activate the cell identity and cell signaling properties to undergo sustained proliferation whereas the CSD cells only undergo early proliferation, and little proliferation at later time points.

**Figure 3.**
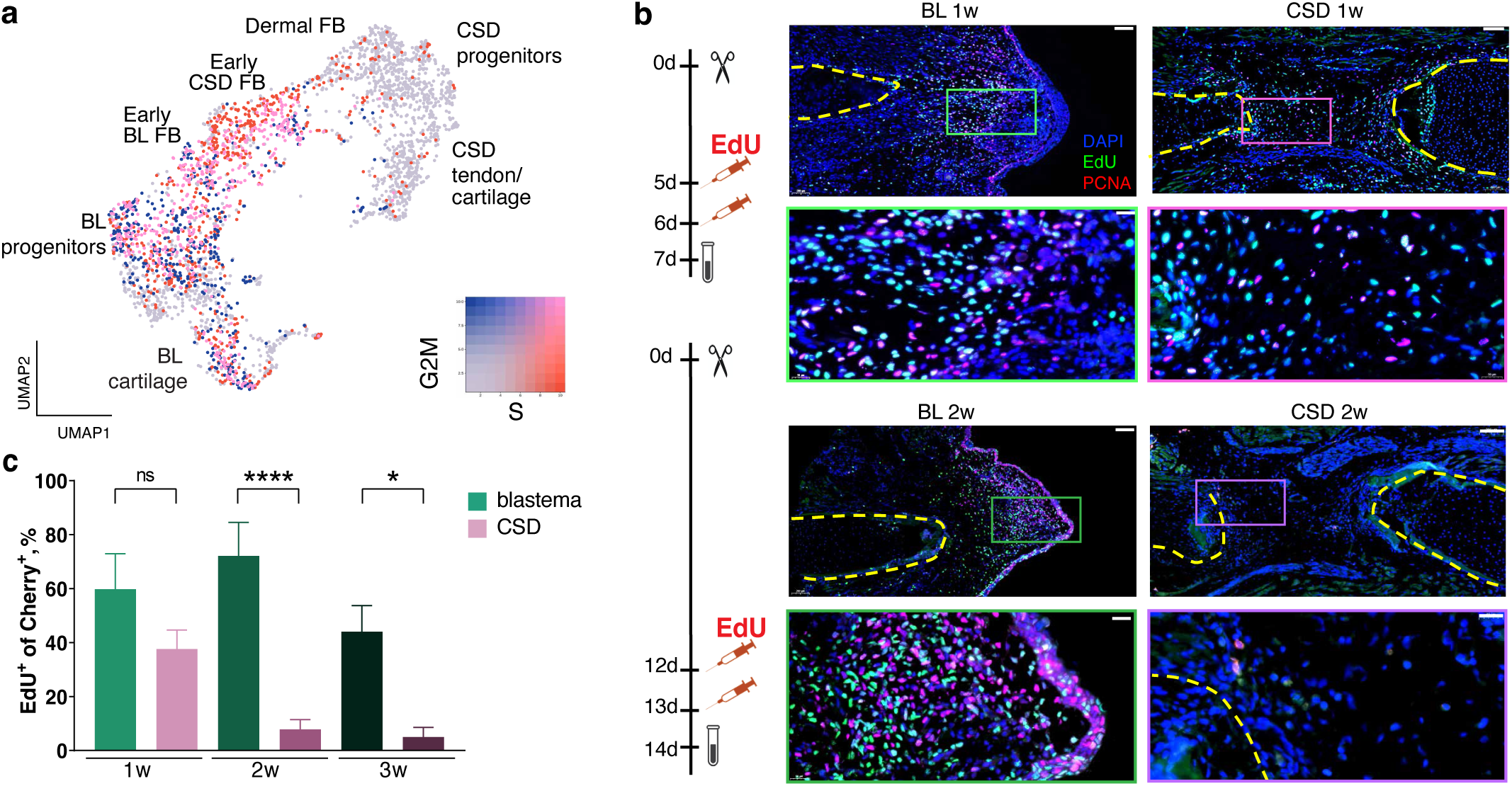
Proliferation is sustained in blastema but diminishes in CSD. **A.** Proliferation score in blastema and CSD-derived connective tissue cells. S-phase score in red and G2M score in blue, the color shade indicates the degree of proliferation score. **B.** EdU injection and sample harvesting schema and representative images. EdU was injected in 2 consecutive days prior to sample harvesting on 1, 2, 3 wpi. In contrast to sustained high proliferation levels in blastema, the initial proliferation rise in CSD at 1-week post-fracture quickly ceased at 2- and 3-weeks post-fracture. Bottom panels are higher magnification images of the framed regions in the upper panels. Blue – DAPI, green – EdU, red – PCNA. Scale bars in lower magnification images, 200 μm; in higher magnification images, 50 μm. N = 4-14 per condition. **C.** EdU^+^ cell quantification in Cherry^+^ CT cell of Prrx1-LP-Cherry limbs with blastema and CSD. * p < 0.05; ** p < 0.01; *** p < 0.001, **** p < 0.0001 with non-parametric ANOVA and Kruskal–Wallis test.

### GRN analysis implicates divergent Wnt signaling between blastema and CSD

Our analysis indicated that blastema and CSD cell states diverge significantly within one wpi that results in drastically different outcomes in proliferation and differentiation. To determine the gene expression states that may be causal for these differences, we generated a gene regulatory network (GRN) model using TENET ^10^ for the blastema versus the CSD states. The GRNs for blastema and CSD consisted of 989 and 2,524 genes, respectively, of which 728 were present in both networks (Figure 4A, Suppl. Table 2, 3). In addition to the network size comparison, we investigated the topological variations that could influence the behavior of the two GRNs. Mathematical modelling of the GRN topologies identified a key difference between the two injuries. The blastema GRN has a topology characteristic of GRNs, in which a few highly connected transcription factors (TFs) coexist with numerous genes having few connections ^11^. The CSD GRN, on the other hand, does not present this structure, showing a decrease in the number of hubs (Figure S5). Therefore, these results may indicate that TFs that are hubs in the blastema GRN are either missing or have lost their importance in the CSD GRN. Identifying these hubs could help to understand molecular pathways that explain the regenerative differences between BL and CSD CT. To identify which genes might be responsible for the changes in behavior of the two GRNs we created a prioritization pipeline that integrates a GRN centrality metric (betweenness centrality) with differential gene expression (Figure S6). Genes with large differences in centrality and differential expression between blastema and CSD at the earliest time-point (3 dpi) were of particular interest, as we anticipated that modifying the activity of regeneration initiators early after injury has the largest effect. Of the 227 genes that met the filtering criteria, 21 were identified as transcription factors (Figure 4B, Suppl. Table 4). From these, the WNT pathway effectors *Lef1* and *Tcf7l2* caught our attention as the network analysis predicted an opposite centrality despite their known structural and functional similarity ^12^: *Lef1* is a hub in the blastema GRN while *Tcf7l2* is a hub in CSD GRN. The scRNA-seq data and qRT-PCR indicates an injury-specific expression of these TFs, with *Lef1* being present only in blastema and *Tcf7l2* being predominantly expressed in CSD (Figure 4C, Figure SA, B, C, D).

**Figure 4.**
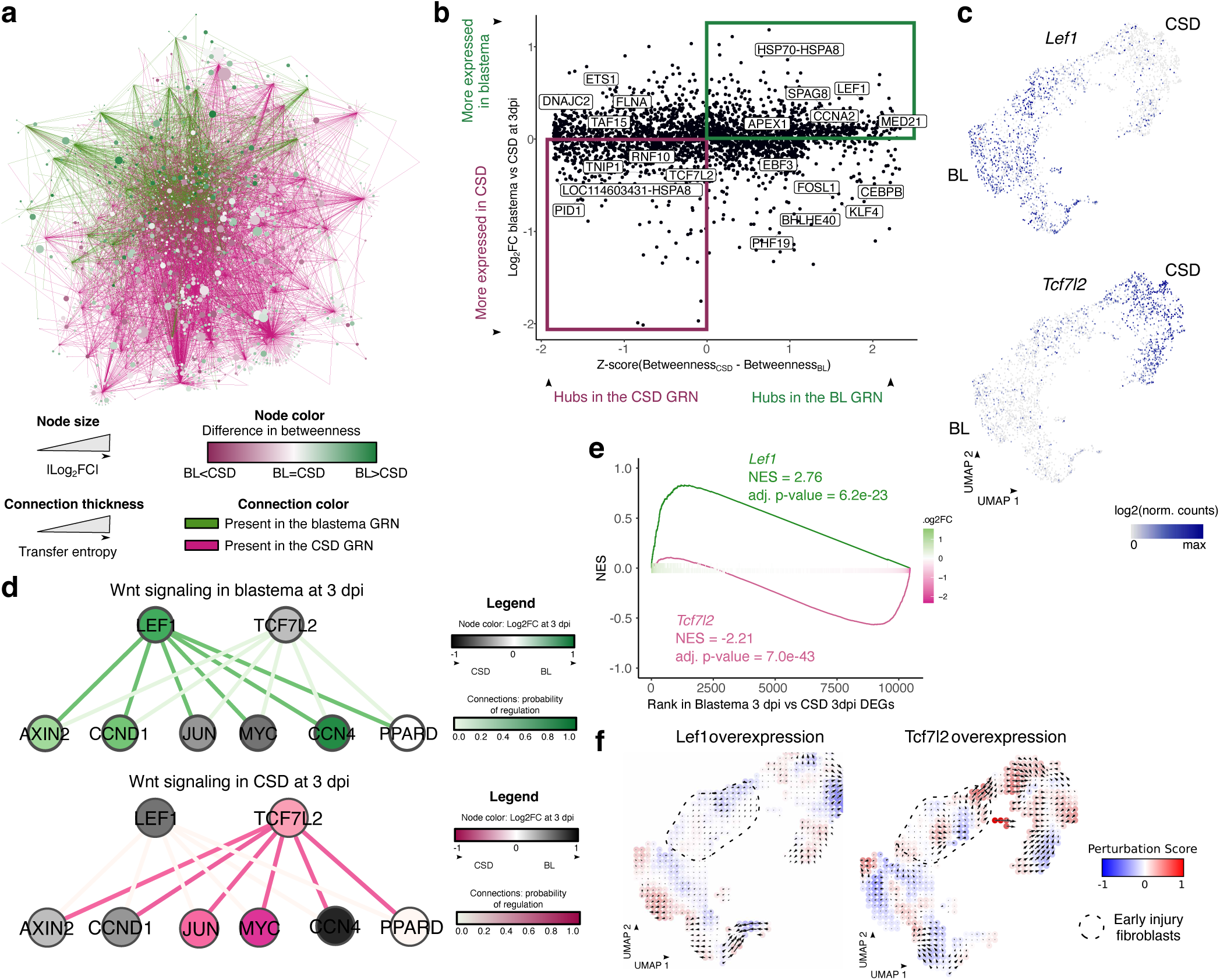
GRN modeling predicts differential Wnt signaling in fibroblasts from blastema and CSD. **a.** Integrated visualization of blastema and CSD CT GRNs. Node size: log2FC between blastema and CSD CT cells at 3 dpi. Node color: difference in ranked betweenness between the blastema and CSD GRNs. Edge color: blastema or CSD CT GRN. Edge thickness: transfer entropy of each link. **b.** Scatterplot of the difference in ranked betweenness between the blastema and CSD CT GRNs expressed as a z-score and the log2FC between blastema and CSD CT at 3 dpi. The names of top candidate TFs are indicated. **c.** UMAP plot of the differential expression of *Lef1* and *Tcf7l2* in blastema and CSD CT. **d.** Predicted regulatory relationships between Lef1 or Tcf7l2 and their target genes in the KEGG Wnt signaling pathway. Node color: log2FC in blastema (green) vs CSD (magenta) at 3 dpi. Connection (edge) color: predicted probability of regulation. **e.** Enrichment plot for the Lef1- and Tcf7l2-specific targets calculated on the log2FC between blastema and CSD at 3 dpi. **f.** Gridded UMAP visualization showing the effect of *Lef1* and *Tcf7l2* overexpression on blastema and CSD CT trajectories, predicted by CellOracle. Red: the perturbation promotes the differentiation trajectory of the cells, blue: the perturbation opposes the differentiation trajectory. Vector arrows indicate the magnitude and direction of the transcriptome changes upon perturbation. The dotted line indicates the early injury fibroblasts.

To further understand how WNT signaling cascades and gene expression programs are differently shaped in blastema and CSD-derived cells, we used a deep learning-based methodology ^13^ trained on 121,098 TF-target gene pairs from axolotl blastema scRNA-seq and bulk ATAC-seq ^14^. Firstly, we investigated whether the two injuries show differences in WNT signaling activation, inhibition and intracellular signaling. The analysis of early (3 dpi) CSD and blastema CT cells revealed differing levels of WNT signaling pathway regulation (Figure 4D, Suppl. Table 5, 6). In terms of transcriptional regulation, the blastema displayed a clear preference for *Lef1*, while CSD demonstrated a strong bias towards *Tcf7l2*. *Lef1* is generally associated with WNT target gene activation, while *Tcf7l2* can act as both an activator and repressor depending on the context ^12^.

Secondly, we used the same algorithm to predict the extent to which *Lef1* and *Tcf7l2* regulate different transcriptional programs. Despite a large overlap in their target gene sets, we found 102 genes uniquely associated with *Lef1* and 1,245 genes uniquely associated with *Tcf7l2* (Figure S7E and Suppl. Table 7, 8). Furthermore, expression of the TF-specific target genes was found to be enriched in blastema and CSD at 3 dpi, respectively (Figure 4E). It is worth noting that *Lef1*-specific targets included matrix metalloproteinases (e.g., *Mmp11*, *Mmp13*) and cell cycle regulators (e.g., *Aurka*, *Cenpf*). In contrast, *Tcf7l2*-specific targets included autophagy modulators (e.g., *Dram2*), Hippo-signaling proteins (e.g., *Sav1*), and WNT inhibitors (e.g., *Dkk1*) (Figure S7F). Altogether, these findings let us hypothesize that *Lef1* and *Tcf7l2* could have antagonistic functions that promote or hinder regeneration, respectively.

To further interrogate the functional effect of these TFs on the CT cell fate, we used the CellOracle algorithm ^15^ to simulate the effect of TF overexpression or KO. The impact of perturbing each TF was estimated by a perturbation score (PS) that runs between −1 (the TF perturbation hinders the differentiation trajectory) and 1 (the TF perturbation pushes the differentiation trajectory). In general, the impact of *Tcf7l2* perturbations was more significant than that of *Lef1* (Figure 4F, S8 A, B, Suppl. Table 9). We focused on the impact of early injury on blastema and CSD fibroblasts, as we hypothesize that early intervention could achieve cellular fate rewiring. The results indicated that the *Lef1* overexpression promoted the blastema trajectory (summarized PS = −0.93, adj. p-value = 5.9e-12), while the *Tcf7l2* overexpression seemed to oppose it (summarized PS = 0.81, adj. p-value = 3.5e-07). As anticipated, the impact of *Lef1* or *Tcf7l2* KO was precisely the inverse of its overexpression (Figure S8 C, D). It is noteworthy that the combined effect of double perturbations was either potentiating or cancelling, depending on the UMAP region (Figure S8).

Based on these findings, we hypothesized that different GRNs could govern blastema and CSD CT identities and may be responsible for their differential behavior and regenerative potential. *Lef1* and *Tcf7l2* may play a significant albeit antagonistic role in both GRNs, which together with their abundance may contribute to the differential regenerative behavior of both injuries.

### Activation of canonical WNT signaling in CSD leads to increased proliferation and CSD bridging with cartilage

Our GRN analysis pointed to *Lef1* activity as a key node specifically associated with the blastema and not the CSD. As *Lef1* is a target gene of canonical WNT signaling ^16, 17, 18^, we pursued the hypothesis that induction of canonical WNT signaling would rescue skeletal formation in the CSD. To induce canonical WNT signaling in the CSD, we encapsulated *in vitro*-transcribed mRNAs of axolotl *Wnt3a* and *Gfp* into Lipid Nanoparticles (LNPs) and injected 0.5 µL LNP suspension into the CSD area at 1, 3, 5, 7, 9, 11 days post-fracture. We first asked if *Wnt3a* expression resulted in prolonged cell proliferation by pulse labeling with EdU prior to tissue harvest at two wpi (Figure 5A). Compared to *Gfp*-injected control CSD, *Wnt3a* expression induced an increase in EdU^+^ cell number in CSD (Figure 5B, C, S9B, C) although the EdU labeling index was lower than observed in the blastema. We then assayed samples at four wpi for formation of bridging cartilage as visualized by alcian blue staining (Figure 5D, Figure S9D). Remarkably, while none of the *Gfp*-expressing control samples showed bridging cartilage, four out of six *Wnt3a*-expressing samples showed bridging cartilage with a gap closure of greater than 90% (Figure 5E, S9D). Given the success of this approach, we also tested several other signaling ligands that were expressed in the early blastema, namely *Inhba* and *Tgfb2* (Figure 5 B-D, S7G-I), which did not induce cell proliferation and bridging cartilage. To understand the underlying transcriptional basis of the *Wnt3a* treatment, we transcriptionally profiled CT cells at 8 dpi from *Wnt3a* versus *Gfp* LNP-treated CSD (Figure 5 F, G). Comparison with CSD and blastema transcriptomes revealed that supplementation with *Wnt3a* resulted in relative down-regulation of several CSD-enriched genes, some of which were previously associated with early phases of regeneration (*Marcks*, *FosB, Fn1*). Blastema-enriched transcripts encoding signaling factors (*Bmp2, Axin2*, *Inhba)* as well as transcripts associated with skeletogenesis (*Bmp2*, *Col24A1*) showed relative up-regulation in the *Wnt3a*-treated sample (Figure 5 G).

**Figure 5.**
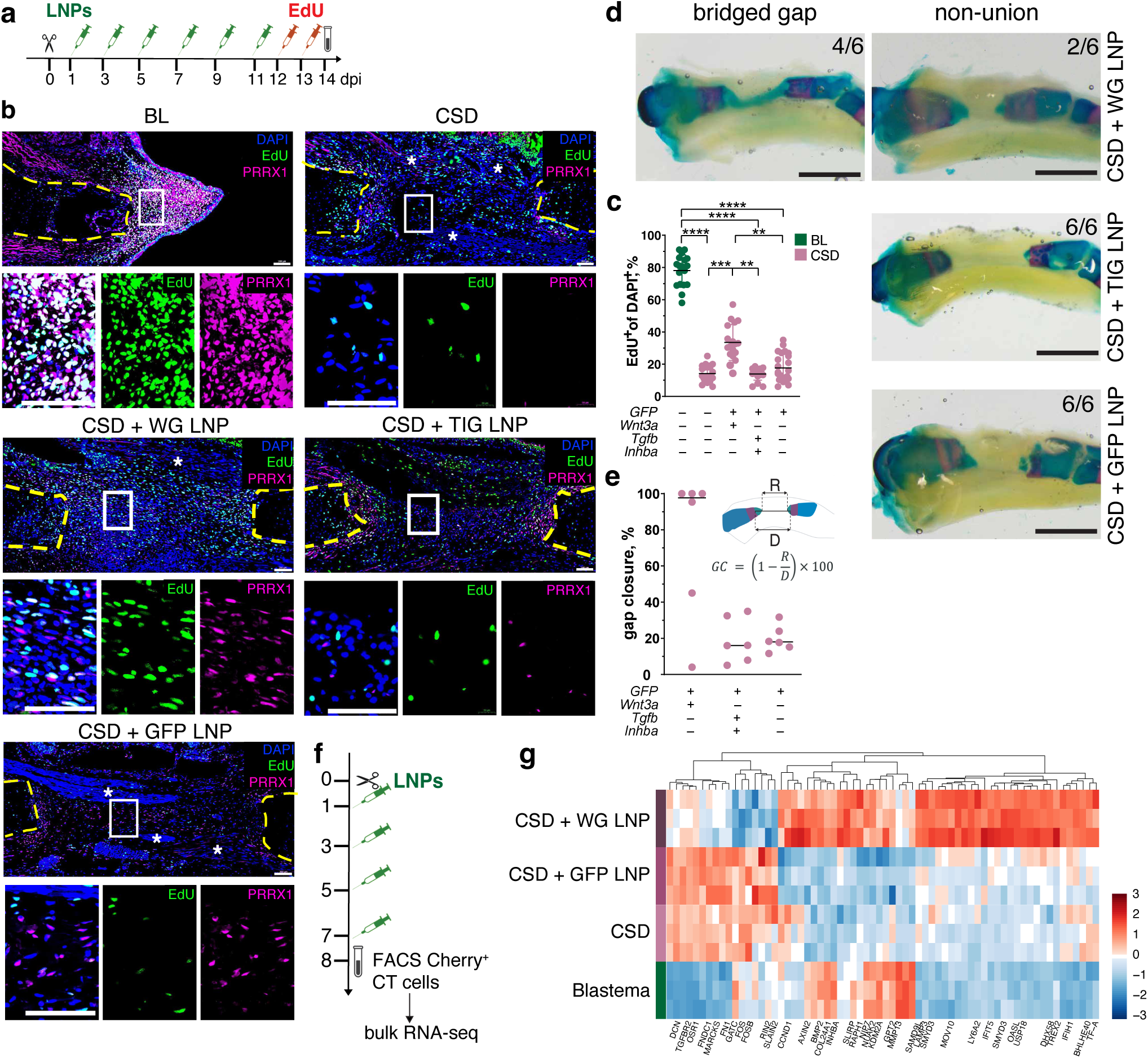
Wnt3a overexpression induces cell proliferation and cartilage formation in the CSD gap. **a.** Schema of the experiment: LNPs were injected at days 1-, 3-, 5-, 7-, 9-, 11-post-fracture and EdU was injected at days 12 and 13 post-injury. **b.** Representative images of EdU and PRRX1 antibody-stained blastema, CSD and CSD with LNP injections at 14 days post-injury. Scale bars 200 µm. **c.** Quantification of EdU-labelled cells in DAPI^+^ cells in blastema, CSD and CSD with LNP injections at 14 days post-injury. Results of 3 independent experiments are presented. * p < 0.05; ** p < 0.01; *** p < 0.001, **** p < 0.0001 with non-parametric multiple comparison one-way ANOVA. N = 17-24 per condition. **d.** Representative images of CSD injected with LNPs and stained with alcian blue/alizarin red 4 weeks post-fracture. LNPs were injected at days 1-, 3-, 5-, 7-, 9-, 11-post-fracture and limbs were harvested at four weeks post-injury. Cartilaginous bridging was observed in 4/6 limbs with *Wnt3a/Gfp* (WG) LNP injections in CSD, and in none of *Tgfb2/Inhba/Gfp* (TIG) and *Gfp* LNP-injected limbs. Scale bar 2 mm. **e.** Gap closure was measured using Ilastik-based segmentation. Horizontal bars show median values. N = 6 per condition. **f.** Schema of the experiment: LNPs were injected at days 1-, 3-, 5-, 7-post-fracture and Cherry^+^ CT cells were FAC-sorted at 8 dpi for bulk RNA-seq analysis. **g.** Hierarchical clustering heatmap of a list of DE genes between CT cells from *Wnt3a/Gfp* LNP and *Gfp* LNP-treated CSD in blastema, CSD, *Wnt3a/Gfp* LNP and *Gfp* LNP-treated CT cells. The color intensity represents normalized and scaled gene expression as LFC, with red indicating high expression and blue indicating low expression. N = 3 in each sample type, p_adj_ < 0.05, FDR < 0.05.

Our studies show that WNT signaling factor is a key signaling modus that is present in the blastema, deficient in the CSD, and is sufficient to induce increased CT proliferation and cartilage bridging of the CSD.

## Discussion

Here we have found a molecular solution to the paradox of the non-regenerating CSD in the highly regenerative axolotl. We demonstrated that in axolotl, blastema precursor CT cells migrate to CSDs and initiate an injury response but fail to proliferate or organize into blastema-like tissue. By mapping the molecular networks that distinguish successful regeneration from persistent defect, we observed reduced activation of WNT signaling pathways. These findings highlight the importance of the local microenvironment in dictating regenerative outcomes. Furthermore, they show that WNT either counteracts the environment hypothesized to prevent bone regeneration, or the environment is not limiting for regeneration in the CSD.

Supplementation of WNT signaling to enhance bone regeneration has been previously reported with some positive effects in improving fracture repair ^20, 21, 22, 23, 24, 25^ but not bridging of CSD as observed here. Bone fracture repair occurs through formation of a callus from bone-associated cell populations such as periosteum that undergo subsequent osteogenesis ^26, 27, 28^, a phenomenon that also occurs at the severed bone ends in axolotl and leads to moderate bone growth^4^. Substantial growth of new skeletal elements during axolotl limb regeneration occurs, however, through the migration of interstitial soft CT cells to the injury site followed by dedifferentiation and recapitulation of skeletal development rather than a post-embryonic bone repair program ^29^. After limb amputation, the wound epidermis provides a source of WNT3a that plays a role in upregulation of FGF8, a key factor involved in sustaining proliferation in mesenchymal blastema cells ^18^. Suppressing WNT signaling via DKK1 overexpression blocks limb regeneration, consistent with an essential role for WNT signaling in blastema cell proliferation and differentiation ^19^. Activation of WNT signaling in the axolotl CSD likely induced the blastema-type program in interstitial CT cells rather enhancement of callus-mediated bone repair. Although the *Wnt3a*-LNPs were injected throughout the CSD without spatial localization, a cartilage rod spanned the CSD anchored to the ends of the remaining bone. How the soft CT in the gap zone organizes and associates with the bone will be fascinating future areas of investigation.

TGF-β signaling has been recognized as pivotal for wound healing, extracellular matrix deposition, but may have a controversial and stage-dependent effect on progenitor proliferation in mammalian fracture healing and chondrogenesis ^30, 31, 32^. As *Tgfb* and *Inhba* were expressed at higher levels in blastema CT cells, we also examined the effects of supplementing these factors in the CSD model which had little effect. TGF-β signaling pathway also regulates cell proliferation and cell fate during tissue development and regeneration. Lack of bone regeneration in the current experiments may be explained by prolonged overexpression of both factors, possibly related to interplay of these two molecules with the other pathways regulating progenitor cell proliferation and fate in the CSD gap or these pathways may function in a different process like signaling to the wound epidermis. Further resolution of application time and doses, as well as combination of *Tgfb* and *Inhba* with canonical WNT can shed light on influence of TGF-β pathway activation on progenitor cells.

By identifying conserved regulatory pathways and intercellular interactions, this work provides a foundation for future interventions aimed at reprogramming non-regenerative environments into regenerative scenarios.

## Methods

### Axolotl husbandry and cell labelling

Axolotls (*Ambystoma mexicanum*) were bred and raised in individual aquaria in the Aquatic facility of the Institute of Molecular pathology, Vienna, Austria. All animal handling and experimental procedures were carried out as approved by the Magistrate of Vienna (GZ: 51072/2019/16, GZ 65248-2021-26), and Federal Ministry of Education, Science and Research (GZ 2024-0.438.711). 6-8 month old, 10-12 cm snout-to-tailtip (snout to the tip of the tail) long axolotls, were used for the fracture surgery and amputations.

To permanently label connective tissue cells with mCherry, axolotl embryos of line tgSceI(Mmu.Prrx1:TFPnls-T2a-Cre-ERT2;CAGGs:LoxP-GFP-GFP-LoxP-Cherry)*^ETNKA^* or tgSceI(Mmu.Prrx1:TFPnls-T2a-Cre-ERT2;CAGGs:LoxP-GFP-dead(STOP)-LoxP-Cherry)*^ETNKA^* were treated with 4-OHT (H7904, Merck) at the forelimb and hindlimb stages as described in Gerber et al., 2018, then raised individually until reached 10-12 cm snout-to-tailtip.

### Bone surgery and amputations in axolotl

The animals were anaesthetized in 0.03% benzocaine solution (Sigma–Aldrich, Germany) for about 15–20 min until good muscle relaxation and lack of reflexive movement upon limb touching with tweezers. Then the animal was placed on a silicone-coated petri dish. A hindlimb was stretched and a polyolefin tube was put on the upper hind limb. The position of the tube was secured using the 7.0 Optilene suture (Braun, Germany). The surgical site was disinfected with 0.75% chlorhexidine. An incision above the femur diaphysis was made using the spring forceps (F.S.T., Germany) and muscles and nerves were carefully displaced from the femur without cutting using fine forceps. The femur was gently lifted by forceps and cut so that about 30% of the bone was removed at the diaphysis area.

For creating blastema samples, the hind limb was amputated at the midshaft of the femur diaphysis using a scalpel blade. To prevent an irregular shape of the blastema, the femur was trimmed to match the level of surrounding tissue.

After the bone fracture and amputation surgery, butorphanol (0.5mg/L) was used as pain medication for 3 days, with daily water change.

### Femur transplantation

The femurs were extracted from 8-9 cm snout-to-tailtip Prrx1-LP-Cherry positive and negative littermate axolotls and transplanted reciprocally maintaining proximal-distal directionality after removing the soft tissues surrounding the bone. The surgical site was disinfected with 0.75% chlorhexidine. Butorphanol (0.5mg/L) was used as pain medication for 3 days post-surgery, with daily water change. 14 days after transplantation, the bone fracture surgery was performed.

### Axolotl genome and transcriptome reference

For sequencing reads mapping and gene annotation, the genome assembly AmexG_v6.0-DD and transcriptome assembly AmexT_v47 were used.

### Isolation of intact soft tissue, CSD and blastema cells for RNA-sequencing and qRT-PCR

For intact soft connective tissue harvesting, upper hind limbs were collected, femurs were removed and cut to 0.5-1 mm pieces prior to enzymatic tissue dissociation. For blastema tissue harvesting, the most distal part of the stump (for samples at 3 dpa and 5 dpa) or complete mesenchymal part of the blastema (for 6, 7, 8, 11 dpa samples) were harvested using razor blade and cut to 0.5-1 mm pieces after peeling off most of the epidermis and removing nerve bundles. For CSD tissue harvesting, the gap tissue excluding skin was collected using 15.0 scalpel (BBraun, 5518032) and large nerve and muscle bundles were removed prior to cutting to 0.5-1 mm pieces. The harvested tissues were dissociated to single cell suspensions using Liberase TM enzyme in 0.7x PBS (Merck, 5401119001) for 50 mins. Then the cells were strained through a 70 μm MACS SmartStrainer (Miltenyi Biotec 130-098-462), sedimented at 300 rcf for 5 mins at 4°C. Cells were resuspended in AMEM (250 ml of MEM w/o L-Glutamine (Gibco, 21090-022), 40 ml of FBS (Sigma-Aldrich, F7524), 4 ml of q mg/mL Insulin (Sigma-Aldrich, 91077C), 4 ml of Penicillin/Streptomycin (Sigma-Aldrich, P0781), 4 ml of Glutamine (Gibco, 25030-024)), filtered with 50 μm Filcon filter (BD Biosciences, 340630) and sorted into 500 µl of TRIzol reagent (Thermo Fisher Scientific, 15596026) or 500 μl MBS (Molecular Biology Service, Vienna BioCenter) RNA isolation kit Lysis buffer with 50mM TCEP (Sigma, 646547). mCherry-positive cell sorting was done with FACSAriaIII^™^ (BD Biosciences) and FACS Diva software v9.0.1 using a 100 μm nozzle and 0.8x FACSFlow^™^ (BD Biosciences). FACS data was re-analyzed using the FLOWJO software (BD Biosciences).

### RNA isolation for RNA-sequencing and qRT-PCR

RNA was extracted according to the TRIzol reagent (Thermo Fisher Scientific, 15596026) manufacturer’s protocol or MBS RNA isolation kit, eluted in 20 μl RNAse-free water and stored at −70°C until required. When the magnetic bead-based RNA extraction protocol (in-house produced) was used, lysis buffer (guanidine thiocyanate, TCEP and Triton X-100-based) containing cells was incubated with paramagnetic beads. The captured RNA was treated with DNase I (New England Biolabs, Germany), washed with consecutive ethanol-based buffer washes and eluted with nuclease-free water.

### Quantitative Reverse Transcription PCR (qRT-PCR)

Complementary DNA was synthesized from 500 ng of total RNA using LunaScript® RT SuperMix (New England Biolabs, Germany). 1µL of reaction mix was used for the qPCR reaction setup, performed with Luna® Universal qPCR Master Mix (New England Biolabs, Germany). Each qRT-PCR reaction was run at least in duplicate. Primers are listed in Suppl. Table 10. Gene expression was normalized to *RP4* (ribosomal protein 4) and the ΔΔCt method^33^ was used to quantify relative fold change in mRNA abundance.

### Library preparation for bulk RNA-sequencing

QuantSeq library preparation and RNA sequencing Libraries for CT cells analysis from LNP-injected samples were prepared using QuantSeq 3′ mRNA-Seq Library Prep Kit FWD (Lexogen) with 2-36 ng of input RNA per sample. Samples were multiplexed for sequencing, using i7+i5 indices 7124:CGTGTGGT+ 5124:CGTGGTTG (CSD with Wnt3a-LNP-1), 7125:GTCCTGGT+ 5125:TGTAGCGG (CSD with Wnt3a-LNP-2), 7126:CCTGCCAA+ 5126:GTGTCATG (CSD with Wnt3a-LNP-3), 7127:TTAATGGT+ 5127:GAACCGGT (CSD control-1), 7128:ATCTACAA+ 5128:CACTCTGA (CSD control-2), 7129:GATATGGT+ 5129:TACACGTT (CSD control-3) 7130:GCCACCAA+ 5130:GAGTGCCT (CSD with GFP-LNP-1), 7131:ATGTTGGT+ 5131:GACGGTCC (CSD with GFP-LNP-2), 7132:GCACAGAC+ 5132:CTGATGCT (CSD with GFP-LNP-3), 7133:ATTGCCTT+ 5133:AATTACCA (Blastema control-1), 7134:GCAATGCA+ 5134:GTCCTGTT (Blastema control-2), 7135:AGTGGCGT+ 5135:GTTAACAG (Blastema control-3). Each replicate was sequenced to a depth of 120 M reads, in PE 150 mode, Illumina NovaSeqX 10B XP flowcell (Illumina). Sequencing was performed by the Next Generation Sequencing Facility at the Vienna BioCenter Core Facilities (VBCF), a member of the Vienna BioCenter (VBC), Austria.

### Data analysis of bulk RNA-sequencing

UMIs were transferred from R1 sequences to read identifiers using the umi2index tool (Lexogen). Additional read processing was performed with BBDuk v38.06-clipping the first 12 nt from the 5’ end (ftl=12) and removing adapter, polyA and low-quality bases from the 3’ end (ref=polyA.fa.gz,truseq.fa.gz k=13 ktrim=r useshortkmers=t mink=5 qtrim=r trimq=10 minlength=20). The processed R1 reads were aligned to the Axolotl genome using hisat2 v2.1.0, PCR duplicates were removed from the alignments using collapse_UMI_bam (Lexogen) and reads in genes were counted with featureCounts (subread v1.6.2) using strand-specific read counting (-s 1). Differential gene expression analysis on raw counts was performed using DESeq2 v1.18.1, with differentially expressed genes identified at an adjusted P-value threshold of padj < 0.05.

### Tissue sections: preparation, staining and imaging

Samples were fixed 24 h in 4% PFA (paraformaldehyde) in PBS, pH 7.4, at 6°C. Then, samples were washed well with PBS and transferred for 48 h to 0.5M EDTA solution for decalcification, at 6°C. After that, samples were washed well with PBS and then equilibrated with 30% sucrose/PBS for 24 h, at 6°C prior to embedding in Tissue-Tek O.C.T. compound (Sakura, Japan), freezing on dry ice and storage at −70°C until sectioning. 12 μm-thick cryosections were prepared using a Cryostar NX70 (Thermo Fisher Scientific) and Epredia MX35 Ultra blades (Thermo Fisher Scientific) and collected on Epredia SuperFrost Plus adhesive microscope slides (Thermo Fisher Scientific). Slides were stored at −20°C until required. For staining, slides were equilibrated to room temperature, then washed well with PBS to remove O.C.T. before proceeding to the following steps.

For Movat’s Pentachrome, tissue was re-hydrated, and incubated for 3 min in 3% acetic acid, for 30 min in 1% Alcian Blue (Sigma Aldrich, Germany)/3% acetic acid, and differentiated for 3-5 min in 3% acetic acid. After washing in ddH2O, sections were incubated for 1 h in ammonia/ethanol, washed twice in tap water, shortly in ddH2O and stained in Weigert’s iron hematoxylin (Chroma-Waldeck, Germany) for 10 min. Sections were washed in tap water and incubated for 15 min in brilliant crocein R fuchsine (Chroma-Waldeck, Germany). After short rinse in 0.5% acidic acid, sections were incubated in 5% phosphotungstic acid (Chroma-Waldeck, Germany) for 20 min and 1 min in 0.5% acetic acid, washed 3x 5 min in 100% EtOH and stained with Saffron-du-Gatinais (Chroma-Waldeck, Germany) for 1h. After dehydrating 3×2 min in 100% EtOH, slides were washed in Xylol and embedded. Staining with Movat’s Pentachrome staining allows for discrimination of fibrous tissue (light green/blue), calcified bone (yellow), muscle (red) and cartilage (intensive green). For Safranin O / Light green staining slides were rehydrated and stained in 1% safranin O (Merck, Germany) for 8 min. Afterwards the sections were dipped in ddH2O 5 times and placed in 0.1% Light green (Chroma-Waldeck, Germany) for 5 min. The sections were then dipped 5 times in 1% acidic acid followed by 2 times for 2 min 100% EtOH treatment. To finish the staining the sections were placed in Xylol 2 times for 2 min and embedded. Safranin O/Light green results in staining bone and soft tissue in green, cartilage-intensive red, and cells in purple/brown.

Prior to PCNA antibody staining, 20 minutes antigen retrieval with DAKO solution (S1699, Agilent, USA) at 80°C was used. For all stainings, the samples were treated for 1 h with the blocking solution (PBS + 0.1% Tween-20 + 5% BSA) at room temperature. The PCNA antibody was applied for 2 h at room temperature, and the rabbit anti-SOX9 (1:500, ZooMAb, MERCK) antibody was applied overnight at 6°C. Slides were washed well with PBS + 0.1% Tween-20 and secondary antibodies conjugated to Alexa fluorophores (Invitrogen) at 1:500 dilution and DAPI at 1:1000 dilution were applied in PBS with 5% BSA with 0,1% Tween-20 for 2 h, RT. Stained slides were washed with PBS + 0.1% Tween-20 and covered with Mowiol/DABCO. The images were captured with a Pannoramic Slide Scanner 250 Flash III (3D HISTECH, Hungary), processed in CaseViewer 2.2 software (3D HISTECH, Hungary) and then colored and merged in FIJI. FIJI software V1.54f/1.8.0. was used for image re-analysis and merging.

### EdU labelling and staining

11 cm snout-to-tailtip axolotls were injected with 10 microgram EdU per gram body weight for 2 days (at 48h and 24h) prior to tissue harvesting. The limbs were fixed in 4% PFA in PBS, pH 7.4 for 24h at 6°C, washed in PBS, de-calcified in 0.5M EDTA for 2 days, 6°C, and treated with 30% sucrose/PBS solution prior to embedding in TissueTek OCT (Science Service, Germany). 12 µm sections were stained with Click-iT EdU Cell Proliferation Kit (ThermoFischer Scientific) according to manufacturer’s protocol and DAPI and scanned as described above.

### HCR in situ hybridization

The 16 μm-thick cryosections were stained according to the HCR RNA-FISH protocol for fixed frozen tissue sections (Molecular Instruments), without the post-fixation and Proteinase K treatment. Probe hybridization buffer and amplification buffer were purchased from Molecular Instruments. HCR probe design was done by identifying unique gene-specific sequences using BLAST alignment against axolotl transcriptome 47. The probe sequences can be found in the Suppl. Table 10. Samples were mounted in Abberior Mount liquid antifade mounting media (Abberior) for imaging. Images were acquired with a LSM980 AxioObserver inverted confocal microscope with ZEN software (Zeiss) with AiryScan2.

### Whole mount alcian blue/alizarin red staining of axolotl limbs

For defining cartilage and calcified bone in axolotl limbs, the limbs were fixed in 4% PFA (Sigma-Aldrich, Germany) in PBS, pH 7.4 for 24h, 6°C and stained with alcian blue/alizarin red. In short, the limbs were washed in PBS, dehydrated in ascending ethanol solutions (25%, 50%, 75%) and then stained in 0.1 mg/mL alcian blue (A3157, Merck) solution. Then, the limbs were rehydrated, washed in 1% KOH solution and subjected to 0.1 mg/mL alizarin red (A5533, Merck) staining. Finally, the limbs were washed with 1% KOH, dehydrated and stored/imaged in 100% glycerol. The imaging was done using an AxioCam ERc 5s color camera (Carl Zeiss Microimaging) mounted on an Olympus SZX10 microscope.

### 10x Genomics experiments and tissue dissociation

Tissue isolation and dissociation were performed as described above. For batch 2, cells were in addition centrifugated in ficoll gradient (25%, 19%, 17%, 15%, 5% in APBS), 30 minutes, 500g, 4°C, washed with APBS, filtered through 30 μm MACS SmartStrainer (BD Biosciences, 130-098-458) and resuspended in APBS + 2% BSA (Bovine Serum Albumin). The cells were loaded into a 10x Genomics microfluidics chip G, encapsulated with barcoded oligo-dT-containing beads using the 10x Genomics Chromium controller according to the manufacturer’s instructions. Single-cell libraries were constructed using a Chromium single-cell 3′ reagent kit (3.1 Chemistry) according to the manufacturer’s instructions. The quality of the cDNA and resulting sequencing libraries were checked by bioanalyzer (High Sensitivity DNA Kit, Agilent, 5067-4626). The libraries were sequenced using Illumina NovaSeq PE150.

### Processing and analysis of scRNAseq data

Sequencing reads were mapped against the axolotl genome ^34, 35^ using STARsolo (STAR 2.7.5a) ^36^ with parameters mimicking Cell Ranger’s (10x Genomics) transcript counting strategy. Fluorescent marker genes were manually added to the references. Seurat v4 ^37^ was mainly used for data analysis. Ribosomal protein genes and pseudo genes (PEG10, L1TD1, RTL, GIN1 and L1-RT) were excluded for downstream analyses. Cells with less than 100 detected genes, a very high (>50,000) or a low (<1,000) number of transcript counts, and cells with a high mitochondrial transcript proportion (10%) were also excluded from the analysis. General processing steps involved Seurat’s default log normalization and scale function to regress out differences in nUMIs. The scaling and the subsequent PCA were always performed using all genes. Uniform Manifold Approximation and Projection (UMAP) was applied to the top 100 principal components (PCs). Data integration was needed as two independent experimental set ups and sequencing runs were performed. We used in all cases Seurat’s built in Harmony function ^38^ to reduce the confounding effect of different batches and samples. Running Harmony was performed with default parameters, 100 input PCs and with selecting the experimental batch as variable to be harmonized in both blastema and CSD, respectively.

Canonical marker genes were used to assign cell type identities to each cluster obtained for blastema and CSD, separately (Suppl. Table 1).

Based on this assignment connective tissue, epidermal, neutrophil and macrophages were extracted and re-clustered, respectively. During the re-clustering process some clusters were identified to be purely driven by low RNA counts and all the cells belonging to these clusters were removed. In addition, the re-clustering revealed additional cells with different identities than aimed for and those cells were removed from the cell type specific data sets.

### CT similarity analysis

After obtaining a cleaned single-cell CT object and removing *Klf5*^+^ cells from the data set we averaged the expression per CT sub cell type using Seurat’s Average Expression function and correlated using Pearson correlation all blastema and CSD subtypes against each other using all genes.

### Trajectory analysis

The diffusion pseudotime presented in Figure S2C was calculated for the CSD and BL samples, respectively, using a diffusion map algorithm, implemented in the R package destiny ^39^ on the harmony matrices using default parameters. This ordered the cells along a pseudo-temporal trajectory that aligns well with the actual sampling time point order. We ranked the cells and normalized the values for each sample, respectively.

### Gene regulatory network inference

Large-scale GRNs were independently reconstructed for blastema and CSD CT cells using TENET (version 2.4), available at https://github.com/neocaleb/TENET ^10^. TENET measures the causal dependency between pairs of genes using transfer entropy (TE), a non-linear, temporally resolved and asymmetric metric based on mutual information ^40^.

For GRN inference, blastema and CSD CT participating in the CT trajectory cells were ordered based on their collection timepoint (3, 5, 6, 7, 8 and 11 dpi). To avoid confounding effects between time and perturbation, the uninjured control (day 0) was not considered for network reconstruction. Next and following TENET recommendations, genes with high expression (log-normalized counts > 1 in 2% of cells) and variability (deviance > quartile 3 for each injury) were selected for GRN reconstruction. Then, TENET was used to calculate the transfer entropy between gene pairs using a default history length (L=1) and potentially indirect relationships in the shape of fast feedforward loops were eliminated if TEx,z < min(TEz,y,TEy,z). Lastly, a statistically significance filter of FDR < 0.01 calculated on one-sided t-tests on standardized TE values was applied to retain statistically significant edges.

### Network topology and power-law analysis

Network topology analysis was performed using Cytoscape v3.9.0 Network Analyzer tool and the igraph R package v1.4.1 in R v4.0.3. Power-law distributions were fit using the igraph R package v1.4.1 and the poweRlaw R package version v0.70.6 in R v4.0.3. Statistical testing of the power-law goodness-of-fit was done using the Kolmogorov-Smirnov test. All network visualizations were done with Cytoscape v3.9.0.

### TF Candidate prioritization

Network topology and gene expression information were integrated to generate a ranking of candidate genes for their perturbation (Figure S6). For each GRN, genes were ranked based on their betweenness centrality:

Betweenness (v) = s≠v≠tds,t(v)ds,t

Where v, s and t refer to nodes in the network and d represents the number of shortest paths between two nodes. Genes absent in one network were added at the end of the ranking with rank N+1, being N the number of genes in that network. Differences in betweenness were calculated by comparing the ranking of each gene in the two GRNs (ΔBetweenness = R(Betweenness)_CSD_ - R(Betweenness)_blastema_, where R(Betweenness) represents ranked betweenness) and standardized. The top 25% genes with largest (positive and negative) standardized differences were pre-selected as candidates.

Gene expression information was incorporated into the prioritization pipeline by calculating the differential gene expression between early injury blastema and CSD CT cells using the Wilcox Rank Summed test in the FindMarkers() function from the Seurat v4 R package ^37^. Genes exhibiting a significant differential expression (Bonferroni corrected p-value < 0.05), an absolute log2FC > than 0.5 and included among the topologically pre-selected candidates were selected for further analysis. Finally, genes annotated as “Transcription factor” or “Cofactor” in AnimalTFDB v4.0 ^41^ were prioritized for further investigation.

### Wnt pathway modeling and prediction of Lef1 and Tcf7l2 target genes

We used Convolutional Neural Network for Co-expression (CNNC), available at https://github.com/xiaoyeye/CNNC ^13^. CNNC was applied in this project to predict gene regulatory relationships between members of WNT signaling pathways and between Lef1/Tcf7l2 and potential target genes.

An axolotl limb-specific convolutional neural network was trained for this project. Training TF-target gene were obtained from Kawaguchi et al., who FACS-sorted CT cells of upper limb 5, 9 and 13 dpi blastemas and profiled their chromatin using by bulk ATAC-seq ^14^. From their identified TF-target gene pairs, those for which both genes have at least 1 log-normalized counts in more than 5% of blastema cells were selected. Negative training examples were generated by randomly selecting pairs of non-interacting genes that pass this expression threshold. A total of 121,098 training cases were used, with a balance between positive and negative training examples. 2-dimensional co-occurrence histograms for the training pairs were built from the blastema 8 and 11 dpi scRNA-seq data generated in this project following the CNNC pipeline. Briefly, normalized gene expression values for each gene were uniformly divided into 32 bins. For each pair of genes, 2-dimensional co-occurrence histograms were generated by assigning each cell to an entry in the matrix and counting the number of samples for each entry. To mitigate the dropout effect, a pseudocount was added to all entries and the matrix was log-transformed. These matrices were visualized as a 32 x 32 histogram and used to train a CNN with the following architecture: one 32 x 32 input layer, 10 intermediate layers (convolutional and maxpooling), 1 flatten layer and a final sigmoid layer. All layers used ReLu as an activation function except the output sigmoid layer. The network was trained using 200 epochs, minibatches of size 1,024 and 3-fold cross-validation.

Once trained, the model was used for two analyses. Firstly, it was used to predict regulatory relationships on 32 x 32 2-dimensional histograms of 3 dpi blastema or CSD scRNA-seq calculated for Lef1 and Tcf7l2 target genes in the “Wnt signaling pathway” as described in the KEGG Pathway Database (entry reference map04310) ^42, 43^, if the genes had at least 1 log-normalized count in at least 5% of blastema or CSD 3 dpi CT cells. Secondly, it was used to predict Lef1 and Tcf7l2 target genes by generating 2-dimensional histograms of the co-expression between Lef1 or Tcf7l2 and all other genes with more than 1 log-normalized count in at least 5% of blastema or CSD 3 dpi CT cells, respectively. In both cases, interactions with a probability > 0.5 were considered positive. Enrichment of Lef1-specific and Tcf7l2-specific target genes in blastema and CSD was calculated with GSEA on the log2FC between blastema and CSD at 3 dpi as implemented in the R package fgsea v1.29.1 ^44^.

### In silico perturbation of candidate TFs

The effect of TF perturbations on the CT transcriptome was predicted using celloracle (version 0.14), available at https://morris-lab.github.io/CellOracle.documentation/ ^15^. For this analysis, the integrated blastema and CSD CT data set and an integrated version of the blastema and CSD CT GRNs inferred with TENET ^10^ containing 2,785 genes and 6,518 edges were used as input. After fitting the linear models, robust connections were identified by filtering Bonferroni-corrected p-values using a significance threshold of 0.05. The effect of a TF perturbation was estimated by propagating the shifts in gene expression through the GRN for 3 iterations. KO simulations assumed absolute TF expression of 0, while overexpression simulations utilized the maximum expression value observed in the dataset. An optimized k value of 81 was used to quantify the effect of the perturbations by correlating the perturbation vector with the gene expression differences between each cell and its 81 k-nearest neighbors. A perturbation score (PS) that estimates the predicted effect of the TF perturbation on the transcriptome was obtained by calculating the inner product between the perturbation vectors and the blastema and CSD trajectories. The PS statistical significance was empirically tested by obtaining a null distribution of the PS calculated with a randomized GRN model. The obtained p-values were Bonferroni-corrected to account for multiple testing.

### Lipid nanoparticles preparation and injection

Lipid nanoparticles, consisting of stabilized target gene(s) mRNA and SM102 (Cayman Chemical)-based lipid mix, were prepared with N/P ratio of 6.0, washed with PBS using Amicon Ultra Centrifugal Filter 10 kDa columns (Merck KgaA, Germany), and diluted with PBS to reach the RNA concentration 0.5µM. The encapsulation efficiency (EE%) was determined with Quant-iT™ RiboGreen RNA Assay Kit (Invitrogen), as previously described^45^. LNPs with EE% above 90% were used for injections with GC120–10 Borosilicate glass capillary (Warner Instruments) and PV830 pneumatic Pico pump (WPI) with ~0.5 µL LNPs injected into the middle of the CSD injury.

### Alcian blue and alizarin red whole-mount staining

For defining cartilage and calcified bone in axolotl limbs, the right fore- and hindlimbs were harvested from 6-6.5 cm snout-to-tailtip axolotl larvae and fixed in 4% PFA in PBS, pH 7.4 for 24 h, +6 °C and stained with alcian blue/alizarin red ^46^. The limbs were washed in PBS, dehydrated in ascending ethanol solutions (25%, 50%, 75%) and then stained in 0.1 mg/mL alcian blue solution. Then, the limbs were rehydrated, washed in 1% KOH solution and subjected to 0.1 mg/mL alizarin red staining. Finally, the limbs were washed with 1% KOH, dehydrated and stored/imaged in glycerol.

### Alcian blue and alizarin red image segmentation and gap size measurement

The raw images were processed in batch with machine-learning based Ilastik ^47^ (v1.4.1) by specifying the labels of old bones and new cartilages. With pixel classification probability in Ilastik, bone masks were identified with probability > 0.7 and cartilages with probability > 0.8. The segmented images were then cropped with a bounding box; then the gap between the bones were measured in the x-axis. Within the gap, the segmented cartilages were checked. If they covered the whole bone gap, then 100% of gap filling was reported; if the gap was partially filled, the length of cartilage was calculated in the x-axis. Finally, the ratios between cartilage length and bone gap were reported.

### Statistical analysis and data representation

Statistical analyses and graph plotting was performed using Prism software (GraphPad, V8). Data were tested for assumptions of normality and equality of variance to determine the appropriate statistical tests to perform. Mean values are reported ± standard deviation (SD). Statistical significance was defined as p_adj_ < 0.05. All figures were assembled in Adobe Illustrator.

## Acknowledgments

We are grateful to Tanaka group, Miguel Torres’ group, the CNIC Bioinformatics Unit members and Dr. Akira Satoh for project discussions. We also thank Yuka Sugiura, Helena Okulski and Pietro Tardivo for laboratory support and Francisco Falcon Chavez and Sergej Nowoshilow for help in bioinformatic analysis. We thank the Animal Care team for axolotl care (Victoria Szilagyi, Andrea Lentz-Koblenc, Erika Kiligan, Magdalena Blaschek, Emina Silic, Elisabeth Zöllner, Tamara Torrecilla Lobos, Dijana Bastian, Julia König, Daniela Pollak and Wilfried Auer). We thank the BioOptics facility and the Molecular Biology Service at the IMP/IMBA Core Facilities for their support. We thank the Next Generation Sequencing facility and Histology facility at the Vienna Biocenter Core Facilities for RNA-sequencing and help with the histological stainings and image acquisition. This research was funded by the ERC (Advanced Grant, 742046 RegGeneMems). A.P. was supported by the Austrian Science Fund (Hertha Firnberg Fellowship number T-1219). I.R.-G received the support of a fellowship from “la Caixa” Foundation (100010434, fellowship code: LCF/BQ/DR20/11790019). M.T. was supported by Comunidad de Madrid grant “CARDIOBOOST” Ref. P2022/BMD-7245; CNIC is supported by the Instituto de Salud Carlos III (ISCIII), the Ministerio de Ciencia e Innovación (MCIN), and the Pro CNIC Foundation, and is a Severo Ochoa Center of Excellence (grant CEX2020-001041-S funded by MICIN/AEI/10.13039/501100011033). F. S.-C. received funding from Severo Ochoa Center of Excellence (grant CEX2020-001041-S funded by MICIN/AEI/10.13039/501100011033) and from the FEDER Una manera de hacer Europa, (grant PID2022-141527OB-I00).

## Author Contributions

A.P., I.R.-G. and T.G. contributed equally to this work. A.P. and E.M.T. conceived the project and secured funding. A.P., E.M.T., F.S.-C., M.T., I.R.-G. and T.G. made substantial contributions to study conception and design. A.P. performed and analyzed the experiments. The bioinformatic analysis was done by T.G., I.R.-G., M.N., J.W. and A.P. with the support from F.C.F. and S.N. A.P., I.R.-G., T.G. and E.M.T. wrote the manuscript. All authors revised and approved the manuscript.

## Data availability

Raw and processed files of the RNA sequencing generated in this study have been deposited under the Gene Expression Omnibus (GEO). The accession number for gene expression data in single-cell RNA-seq experiment is GSE314016, and of bulk RNA-seq of blastema and CSD CT cells is GSE312556. All other formats of raw data are available upon request. Supplementary Information is available for this paper.

## Materials and Correspondence

Correspondence and requests for materials should be addressed to A.P., F.S.-C. and E.M.T.

## SUPPLEMENTARY FIGURES LEGENDS

**Supplementary Figure 1.**
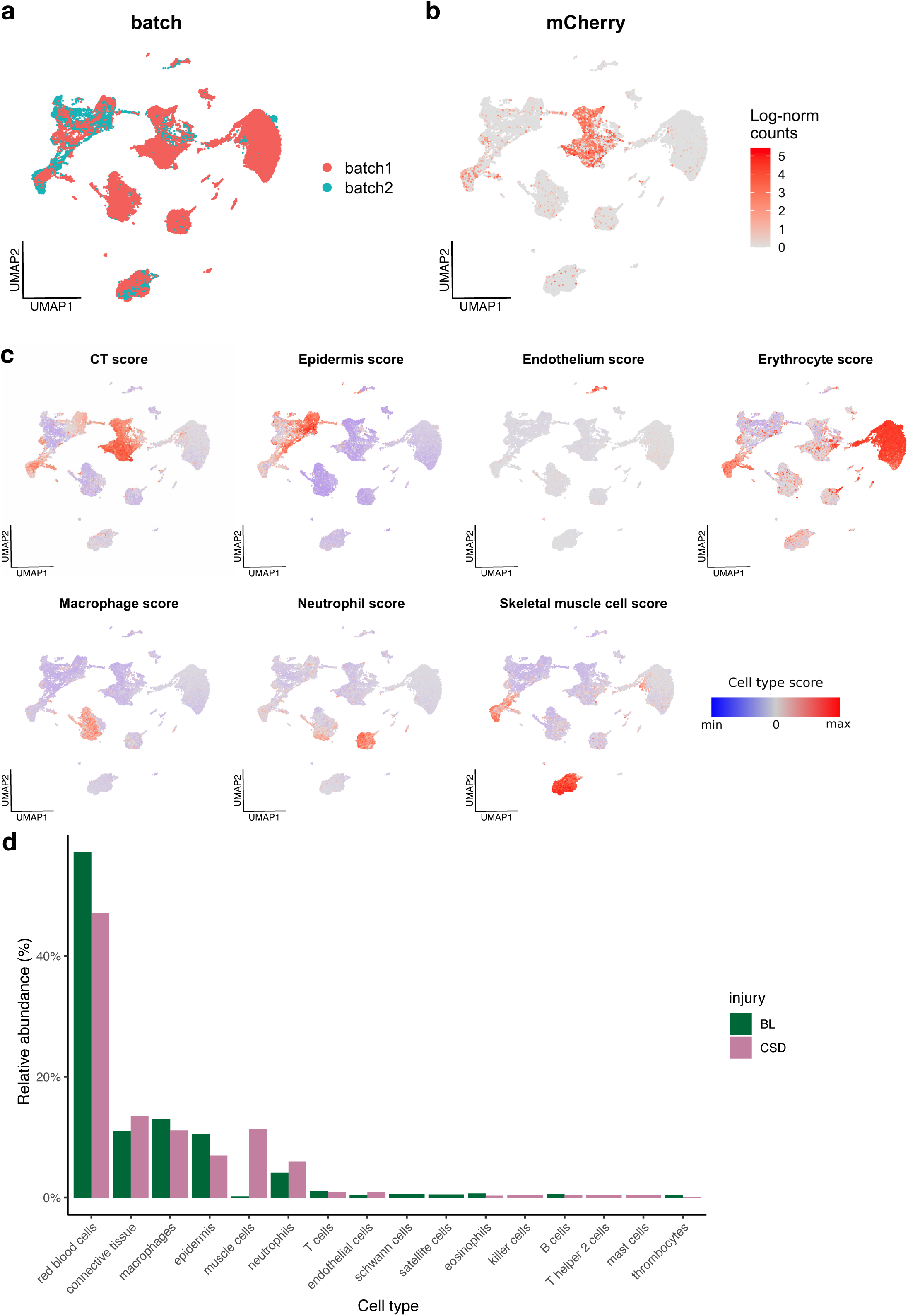
Cellular composition of blastema and CSD single-cell transcriptomics samples. **a.** UMAP of integrated blastema and CSD cells colored by batch. **b.** UMAP of integrated blastema and CSD cells colored by log-normalized mCherry expression. **c**. UMAPs of integrated blastema and CSD cells colored by cell type scores calculated from the expression of known cell type marker genes. **d**. Relative abundance of cell types in blastema and CSD samples.

**Supplementary Figure 2.**
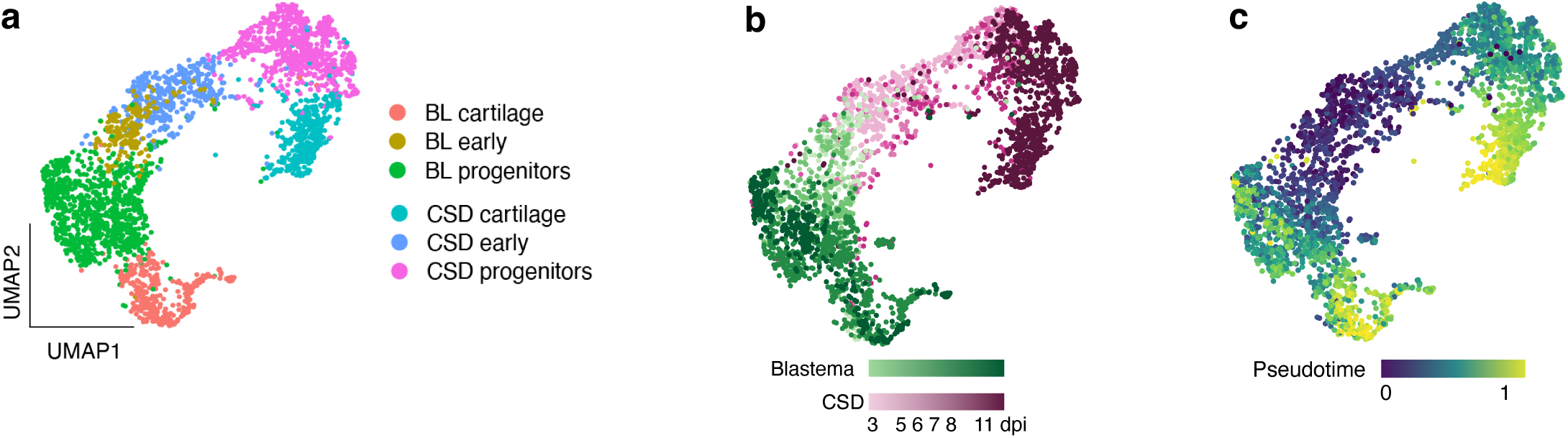
Trajectory inference for blastema and CSD CT. **a.** UMAP of integrated blastema and CSD CT cells colored by cluster. **b**. UMAP of integrated blastema and CSD CT cells colored by injury and time-point. **c.** UMAP of integrated blastema and CSD CT cells colored by pseudotime estimates.

**Supplementary Figure 3.**
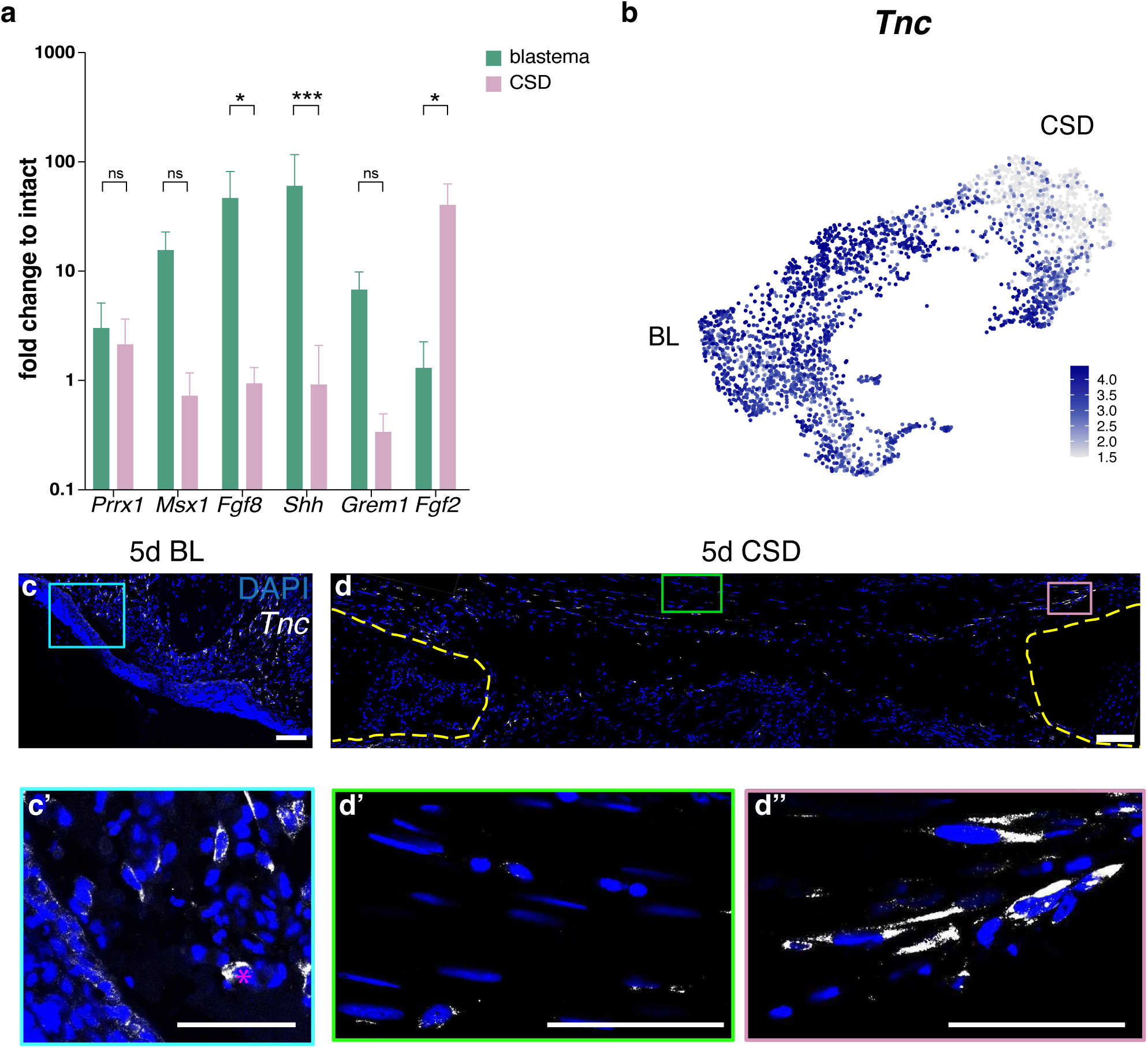
Validation of differentially expressed genes in blastema and CSD. **a.** qRT-PCR validation of the DE genes in sorted blastema and CSD CT cells 8 days post-injury (corresponding to mid-stage blastema). Data was normalized to the expression in intact cells. Note lower expression of the mid-stage blastema genes and upregulation of early blastema gene, *Fgf2*, in CSD CT cells. n = 4-6. * p < 0.05; ** p < 0.01; *** p < 0.001 with 2-way ANOVA test. **b.** UMAP plot of the expression of *Tnc* in blastema and CSD CT cells. **c, d.** HCR *in situ* hybridization was used to visualize and localize the *Tnc* transcripts (white dots) in blastema and CSD 5 days post-injury. Note high *Tnc* expression in the blastema cells, as well as at the CSD edges and a few mid-CSD cells. Scale bars: c, d - 200 µm, c’, d’, d’’ - 100 µm. Magenta asterisks mark autofluorescence.

**Supplementary Figure 4.**
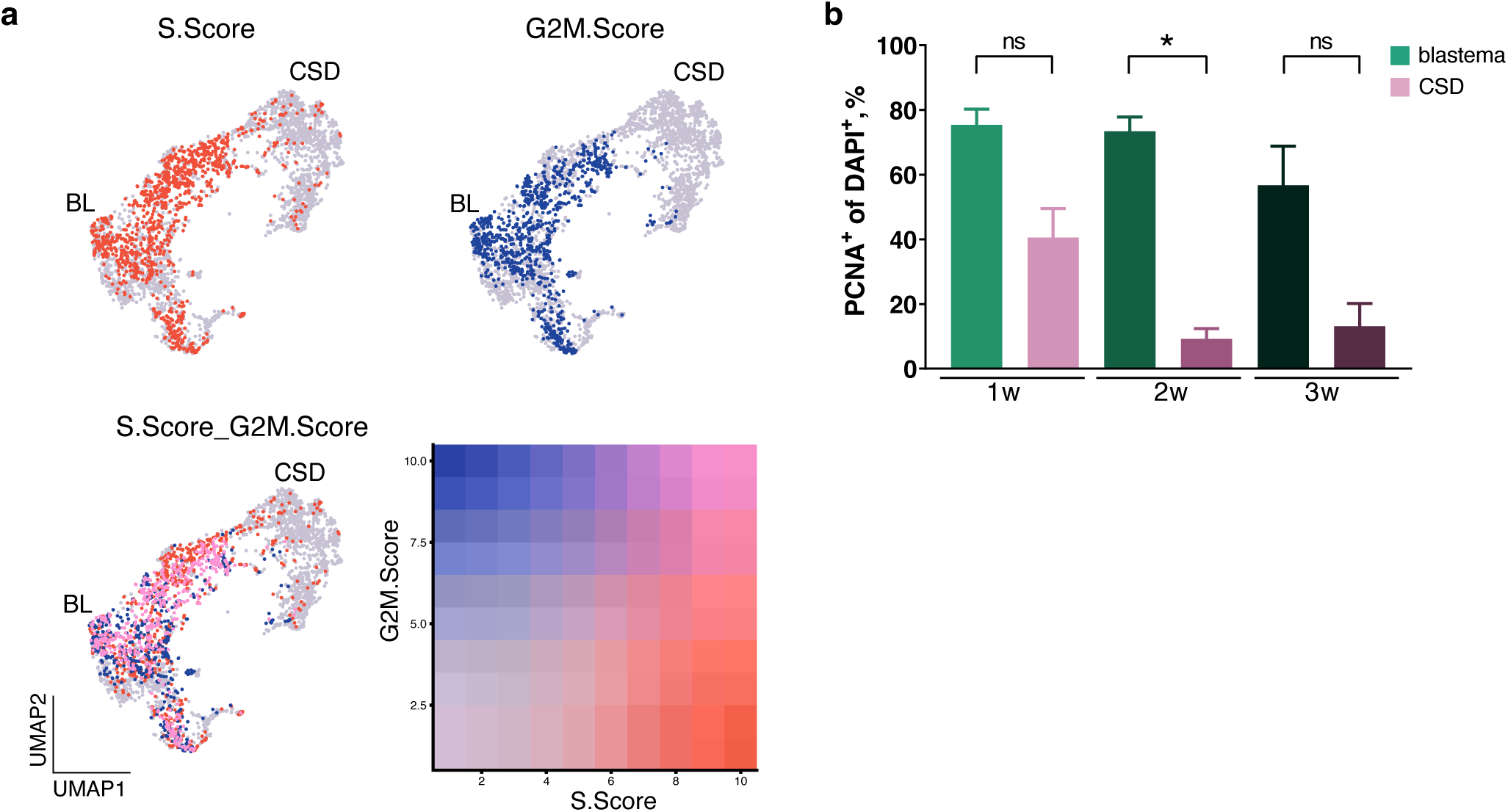
Proliferation dynamics in blastema and CSD. **a.** Proliferation score in blastema and CSD-derived connective tissue cells. S-phase score in red and G2M score in blue, the color shade indicates the degree of proliferation score. **b.** Quantification of PCNA-positive cells in blastema and CSD 1-, 2- and 3-weeks post-injury. * p < 0.05 with non-parametric ANOVA. N = 4 per condition/ time point.

**Supplementary Figure 5.**
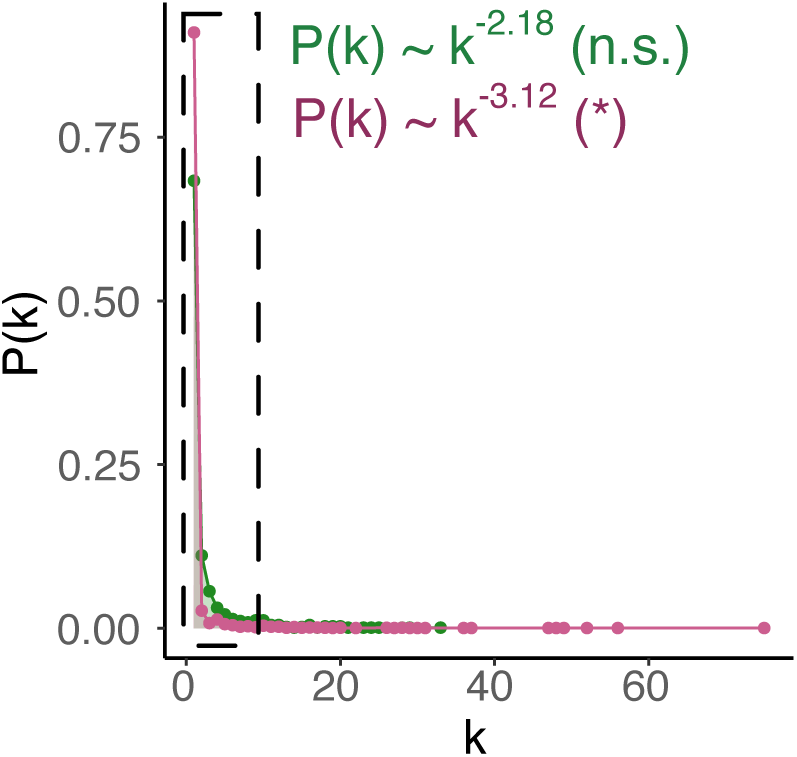
Power-law fit of blastema (green) and CSD (magenta) CT GRN out-degree. Most GRN have been described to follow a scale-free and hierarchical topology in which a few highly connected “hub” nodes coexist with numerous nodes having relatively few connections. Scale-free networks can be easily identified because their degree distribution (probability that a chosen node has exactly k edges) follows a power law P(k) ~k-γ, where 2<γ<3 ^11^. The out-degree distribution for the blastema GRN followed a power law with a degree exponent of Blastema = 2.17 (p-value = 0.79). However, the out-degree distribution for the CSD GRN could not be adequately described by a power law (CSD=3.12 and p-value = 3.9e-9). The p-value corresponds to a Kolmogórov-Smirnov goodness-of-fit test. A p-value > 0.05 (n.s.) means that the observed distribution fits the power-law distribution, while a p-value < 0.05 (*) means that the observed distribution cannot be properly described by a power-law distribution.

**Supplementary Figure 6.**
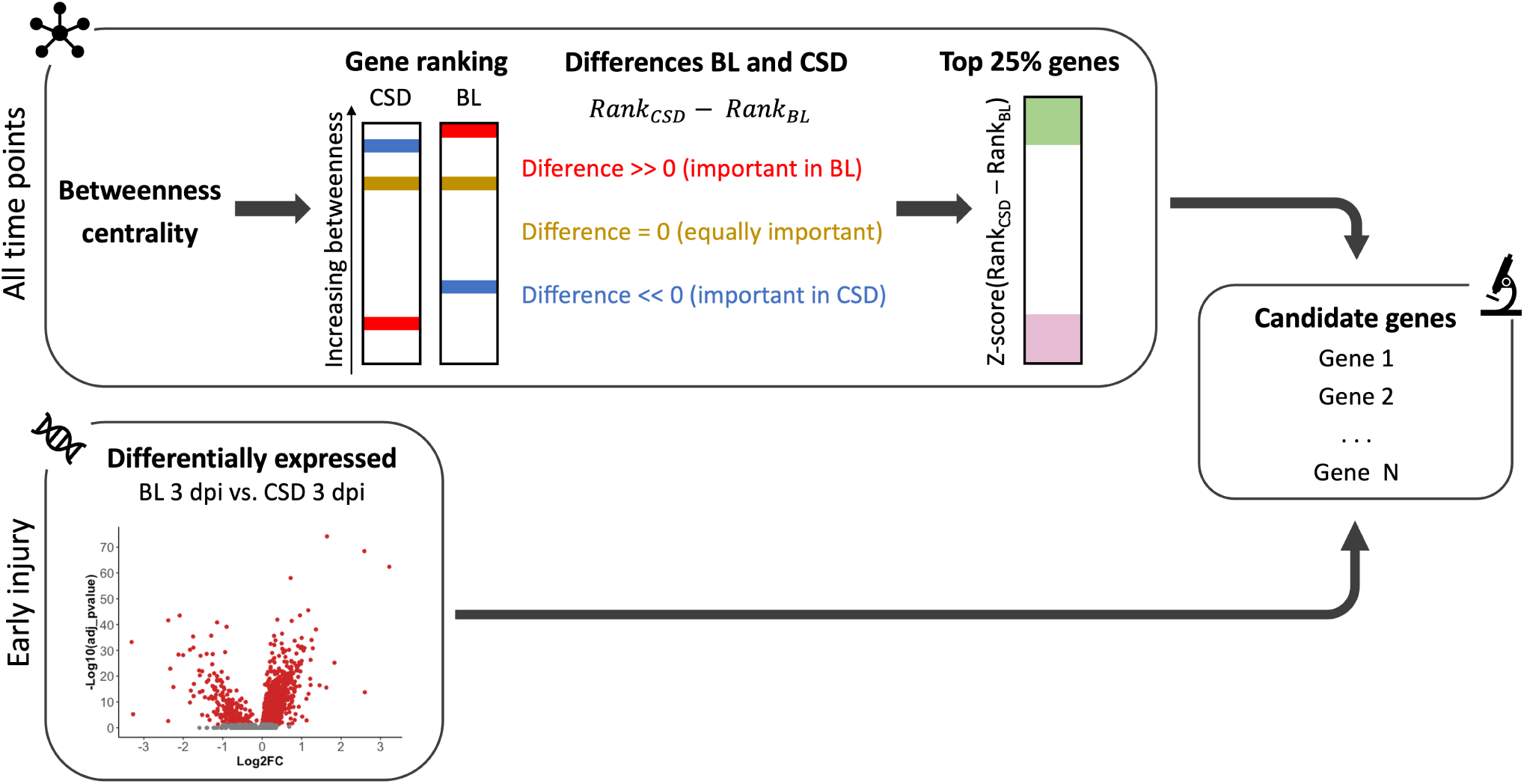
Pipeline for candidate selection. Genes were independently ranked by betweenness centrality in the blastema and CSD GRNs. Genes absent in a network received the ranking N + 1, with N being the number of genes in the network. The two rankings were compared to find the genes with the largest difference in relative betweenness between the two GRNs. Top 25% genes for each network were pre-selected as candidates. This list was pruned to keep only those genes with a significant differential expression (Bonferroni-adjusted p-value < 0.05) and absolute log2FC > 0.5 between blastema and CSD at 3 dpi.

**Supplementary Figure 7.**
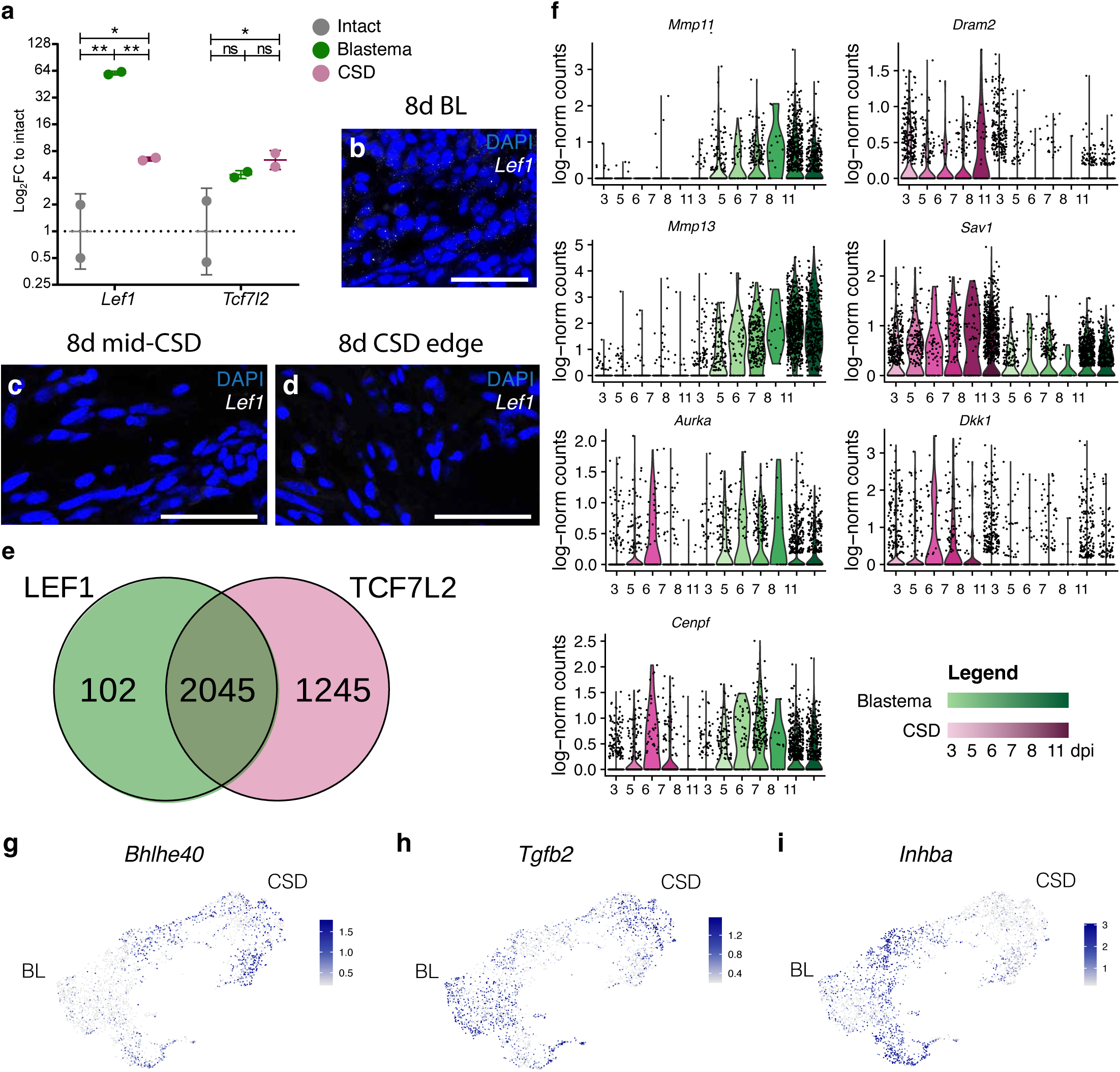
Expression of Lef1, Tcf7l2, their target genes and TGF-β pathway genes in blastema and CSD. **a.** qRT-PCR of *Lef1* and *Tcf7l2* in intact CT (0 dpi), blastema CT (11 dpi) and CSD CT (11 dpi). * p < 0.05; ** p < 0.0001, ns – non-significant with 2-way ANOVA test with Turkey’s multiple comparison tests. **b, c, d.** HCR in situ hybridization was used to visualize and localize the *Lef1* transcripts (white dots) in blastema and CSD 8 days post-injury. Note *Lef1* expression in the blastema epidermis and adjacent CT, and lower expression in CSD cells around the fracture edges and at the mid-CSD gap. Scale bars - 100 µm. **e.** Venn diagram showing the number of shared and unique genes associated with *Lef1* and *Tcf7l2*. **f.** Violin plots showing the changes in gene expression in the two injuries over time for some *Lef1*-specific target genes (*Mmp11*, *Mmp13*, *Aurka*, and Cenpf) and for some *Tcf7l2*-specific target genes (*Dram2*, *Sav1*, *Dkk1* and *Gas6*). **g.** UMAP plot of the expression of *Bhlhe40* in blastema and CSD CT cells. **h.** UMAP plot of the expression of *Tgfb2* in blastema and CSD CT cells. **i.** UMAP plot of the expression of *Inhba* in blastema and CSD CT cells.

**Supplementary Figure 8.**
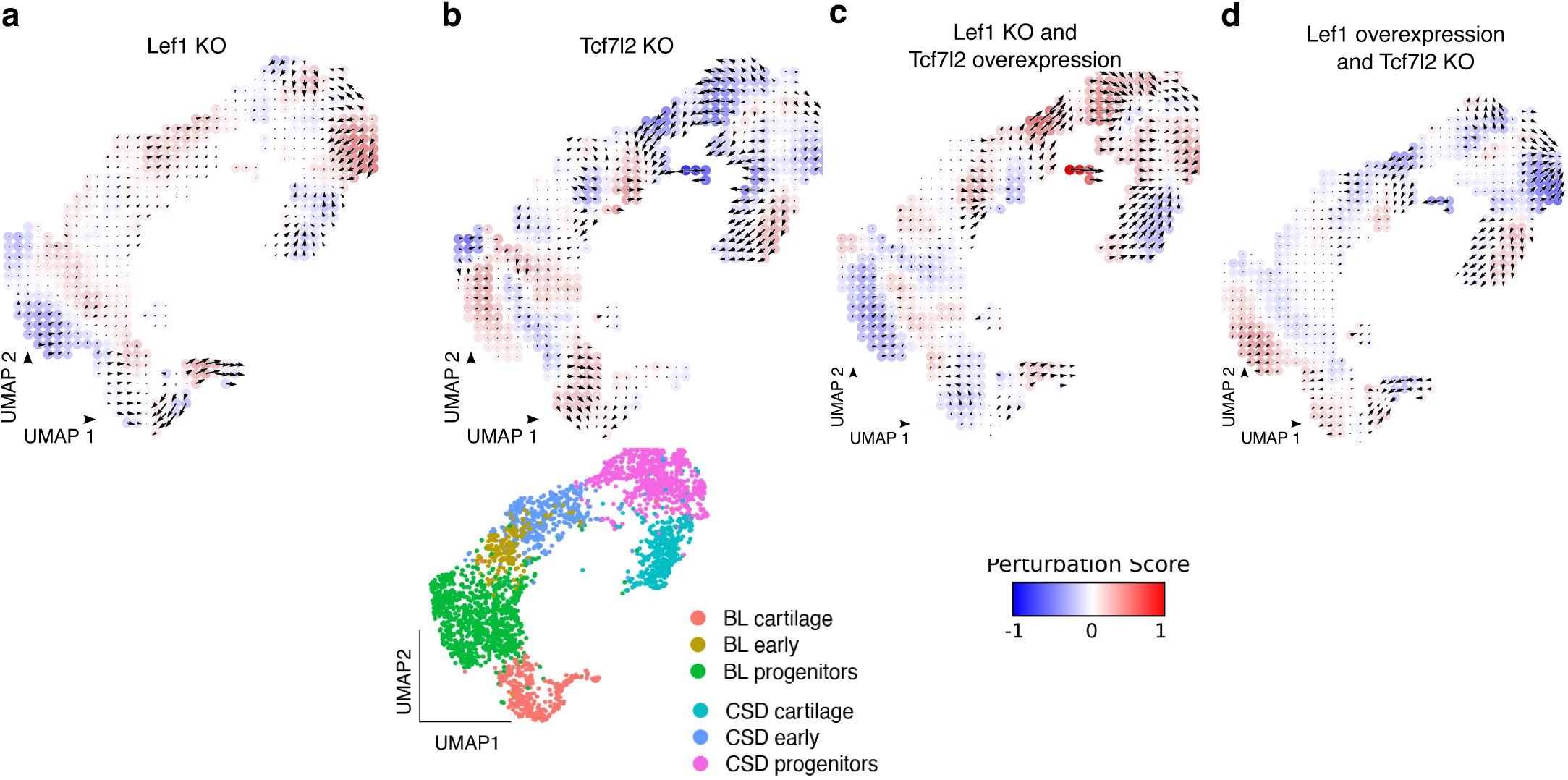
Effect of Lef1 and Tcf7l2 in silico perturbations on the blastema and CSD CT trajectories. Gridded UMAP visualization of integrated blastema and CSD CT cells. Colors and vectors represent the effect of **a**. *Lef1* knock-out, **b**. *Tcf7l2* knock-out, **c**. simultaneous Lef1 KO and *Tcf7l2* overexpression and **d**. simultaneous *Lef1* overexpression and *Tcf7l2* KO. Red colors indicate that the perturbation promotes the differentiation trajectory of the cells, while blue colors indicate that the perturbation opposes the differentiation trajectory. Vector arrows indicate the magnitude and direction of the transcriptome changes upon perturbation.

**Supplementary Figure 9.**
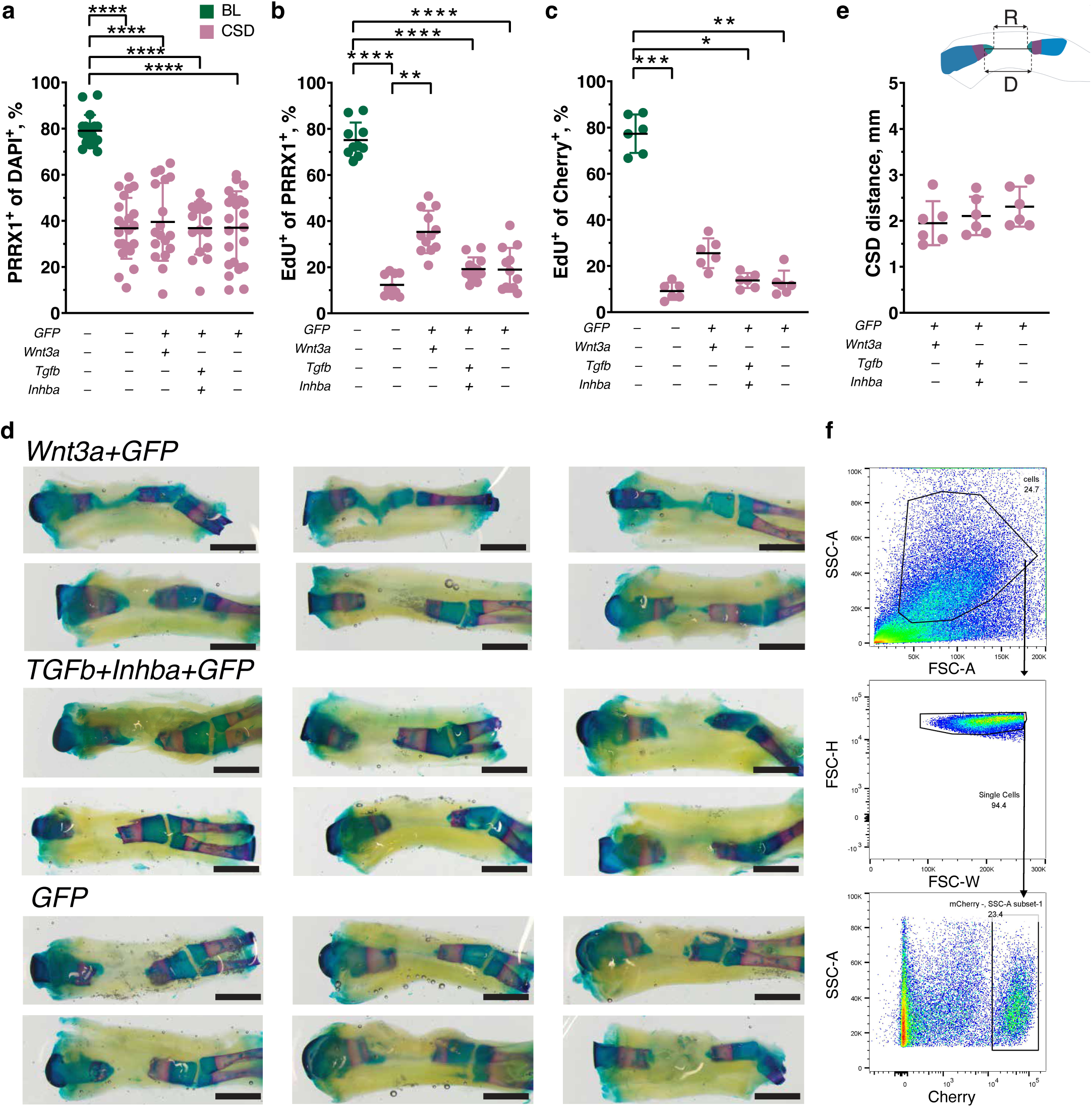
Effect of WNT and TGF-β pathway perturbations on cell proliferation and bone regeneration in CSD. **a**. Quantification of PRRX1 antibody-labelled cells in blastema, control CSD and CSD samples injected with LNPs. Results of 2 independent experiments are presented. * p < 0.05; ** p < 0.01; *** p < 0.001, **** p < 0.0001 with non-parametric multiple comparison one-way ANOVA. **b**. Quantification of EdU-labelled cells among PRRX1 antibody-stained cells in blastema, control CSD and CSD samples injected with LNPs. Results of 2 independent experiments are presented. * p < 0.05; ** p < 0.01; *** p < 0.001, **** p < 0.0001 with non-parametric multiple comparison one-way ANOVA. **c**. Quantification of EdU-labelled cells among mCherry^+^ cells in *Prrx1-LP-Cherry* axolotls with blastema, control CSD and CSD samples injected with LNPs. Results of 2 independent experiments are presented. * p < 0.05; ** p < 0.01; *** p < 0.001, **** p < 0.0001 with non-parametric multiple comparison one-way ANOVA. N = 6. **d.** CSD injected with LNPs and stained with alcian blue/alizarin red 4 weeks post-fracture. LNPs were injected at days 1-, 3-, 5-, 7-, 9-, 11-post-fracture and limbs were harvested at four weeks post-injury. Cartilaginous bridging was observed in 4/6 limbs with *Wnt3a/Gfp* LNP injections in CSD, and in none of *Tgfb2/Inhba/Gfp* and *Gfp* LNP-injected limbs. Scale bars 2 mm. **e**. Measurement of the CSD gap in the CSD samples injected with LNPs, 4 weeks post-fracture. **f**. Gating strategy for flow cytometry of Cherry^+^ cells in *Prrx1-LP-Cherry* axolotls. Representative plots depicting gates defined for all cells (upper panel), single cells (centre) and Cherry-positive cells (bottom). Depicted are plots from the 8 dpi CSD sample.

**Supplementary Table 1.** Marker table for cells.

**Supplementary Table 2.** Blastema GRN.

**Supplementary Table 3.** CSD GRN.

**Supplementary Table 4.** Candidate prioritization results.

**Supplementary Table 5.** Wnt signaling network for blastema.

**Supplementary Table 6.** Wnt signaling network for CSD.

**Supplementary Table 7.** Predicted *Lef1* target genes in blastema.

**Supplementary Table 8.** Predicted *Tcf7l2* target genes in CSD.

**Supplementary Table 9.** Results of perturbations with CellOracle.

**Supplementary Table 10.** qPCR primers and HCR probes sequences.

**Supplementary Table 11.** DEseq2 results of genes expressed in CSD with *GFP* LNP vs CSD with *Wnt3a* LNP.

## References

1. Chen, X., Song, F., Jhamb, D., Li, J., Bottino, M. C., Palakal, M. J. & Stocum, D. L. The axolotl fibula as a model for the induction of regeneration across large segment defects in long bones of the extremities. PLoS One 10, e0130819 (2015). 10.1371/journal.pone.0130819

2. Satoh, A., Cummings, G. M., Bryant, S. V. & Gardiner, D. M. Neurotrophic regulation of fibroblast dedifferentiation during limb skeletal regeneration in the axolotl (*Ambystoma mexicanum*). Dev. Biol. 337, 444–457 (2010). 10.1016/j.ydbio.2009.11.023

3. Currie, J. D., Kawaguchi, A., Traspas, R. M., Schuez, M., Chara, O. & Tanaka, E. M. Live imaging of axolotl digit regeneration reveals spatiotemporal choreography of diverse connective tissue progenitor pools. Dev. Cell 39, 411–423 (2016). 10.1016/j.devcel.2016.10.013

4. Gerber, T., Murawala, P., Knapp, D., Masselink, W., Schuez, M., Hermann, S., Gac-Santel, M., Nowoshilow, S., Kageyama, J., Khattak, S., Currie, J. D., Camp, J. G., Tanaka, E. M. & Treutlein, B. Single-cell analysis uncovers convergence of cell identities during axolotl limb regeneration. Science 362, eaaq0681 (2018). 10.1126/science.aaq0681

5. Khattak, S., Schuez, M., Richter, T., Knapp, D., Haigo, S. L., Sandoval-Guzmán, T., Hradlikova, K., Duemmler, A., Kerney, R. & Tanaka, E. M. Germline transgenic methods for tracking cells and testing gene function during regeneration in the axolotl. Stem Cell Reports 1, 90–103 (2013). 10.1016/j.stemcr.2013.03.002

6. Bryant, D. M., Johnson, K., DiTommaso, T., Tickle, T., Couger, M. B., Payzin-Dogru, D., Lee, T. J., Leigh, N. D., Kuo, T. H., Davis, F. G., Bateman, J., Bryant, S., Guzikowski, A. R., Tsai, S. L., Coyne, S., Ye, W. W., Freeman, R. M. Jr, Peshkin, L., Tabin, C. J., Regev, A., Haas, B. J. & Whited, J. L. A tissue-mapped axolotl de novo transcriptome enables identification of limb regeneration factors. Cell Rep. 18, 762–776 (2017). 10.1016/j.celrep.2016.12.063

7. Tsujioka, H., Kunieda, T., Katou, Y., Shirahige, K., Fukazawa, T. & Kubo, T. Interleukin-11 induces and maintains progenitors of different cell lineages during Xenopus tadpole tail regeneration. Nat. Commun. 8, 495 (2017). 10.1038/s41467-017-00594-5

8. García-García, D., Knapp, D., Kim, M., Jamwal, K., Fuqua, H., Seaman, R. P., Grindle, R. E., Nowoshilow, S., Novatchkova, M., Kolling, F. W., Graber, J. H. & Murawala, P. The essential role of connective-tissue cells during axolotl limb regeneration. bioRxiv 2025.03.30.645595 (2025). 10.1101/2025.03.30.645595

9. Nacu, E., Gromberg, E., Oliveira, C. R., Drechsel, D. & Tanaka, E. M. FGF8 and SHH substitute for anterior-posterior tissue interactions to induce limb regeneration. Nature 533, 407–410 (2016). 10.1038/nature17972

10. Kim, J., Jakobsen, S. T., Natarajan, K. N. & Won, K. J. TENET: gene network reconstruction using transfer entropy reveals key regulatory factors from single cell transcriptomic data. Nucleic Acids Res. 49, e1 (2021). 10.1093/nar/gkaa1014

11. Barabási, A. L. & Oltvai, Z. N. Network biology: understanding the cell’s functional organization. Nat. Rev. Genet. 5, 101–113 (2004). 10.1038/nrg1272

12. Cadigan, K. M. & Waterman, M. L. TCF/LEFs and Wnt signaling in the nucleus. Cold Spring Harb. Perspect. Biol. 4, a007906 (2012). 10.1101/cshperspect.a007906

13. Yuan, Y. & Bar-Joseph, Z. Deep learning for inferring gene relationships from single-cell expression data. Proc. Natl Acad. Sci. USA 116, 27151–27158 (2019). 10.1073/pnas.1911536116

14. Kawaguchi, A., Wang, J., Knapp, D., Murawala, P., Nowoshilow, S., Masselink, W., Taniguchi-Sugiura, Y., Fei, J. F. & Tanaka, E. M. A chromatin code for limb segment identity in axolotl limb regeneration. Dev. Cell 59, 2239–2253.e9 (2024). 10.1016/j.devcel.2024.05.002

15. Kamimoto, K., Stringa, B., Hoffmann, C. M., Jindal, K., Solnica-Krezel, L. & Morris, S. A. Dissecting cell identity via network inference and in silico gene perturbation. Nature 614, 742–751 (2023). 10.1038/s41586-022-05688-9

16. Galceran, J., Fariñas, I., Depew, M. J., Clevers, H. & Grosschedl, R. Wnt3a-/--like phenotype and limb deficiency in Lef1(-/-)Tcf1(-/-) mice. Genes Dev. 13, 709–717 (1999). 10.1101/gad.13.6.709

17. Poss, K. D., Shen, J. & Keating, M. T. Induction of lef1 during zebrafish fin regeneration. Dev. Dyn. 219, 282–286 (2000). 10.1002/1097-0177(2000)9999:9999<::aid-dvdy1045>3.3.co;2-3

18. Glotzer, G. L., Tardivo, P. & Tanaka, E. M. Canonical Wnt signaling and the regulation of divergent mesenchymal Fgf8 expression in axolotl limb development and regeneration. eLife 11, e79762 (2022). 10.7554/eLife.79762

19. Kawakami, Y., Rodriguez Esteban, C., Raya, M., Kawakami, H., Martí, M., Dubova, I. & Izpisúa Belmonte, J. C. Wnt/beta-catenin signaling regulates vertebrate limb regeneration. Genes Dev. 20, 3232–3237 (2006). 10.1101/gad.1475106

20. Nelson, A. L., Mancino, C., Gao, X., Choe, J. A., Chubb, L., Williams, K., Czachor, M., Marcucio, R., Taraballi, F., Cooke, J. P., Huard, J., Bahney, C. & Ehrhart, N. β-catenin mRNA encapsulated in SM-102 lipid nanoparticles enhances bone formation in a murine tibia fracture repair model. Bioact. Mater. 39, 273–286 (2024). 10.1016/j.bioactmat.2024.05.020

21. Haffner-Luntzer, M., Ragipoglu, D., Ahmad, M., Schoppa, A., Steppe, L., Fischer, V., Luther, J., Yorgan, T., Bockamp, E., Amling, M., Schinke, T. & Ignatius, A. Wnt1 boosts fracture healing by enhancing bone formation in the fracture callus. J. Bone Miner. Res. 38, 749–764 (2023). 10.1002/jbmr.4797

22. Chen, Y., Whetstone, H. C., Lin, A. C., Nadesan, P., Wei, Q., Poon, R. & Alman, B. A. Beta-catenin signaling plays a disparate role in different phases of fracture repair: implications for therapy to improve bone healing. PLoS Med. 4, e249 (2007). 10.1371/journal.pmed.0040249

23. Minear, S., Leucht, P., Jiang, J., Liu, B., Zeng, A., Fuerer, C., Nusse, R. & Helms, J. A. Wnt proteins promote bone regeneration. Sci. Transl. Med. 2, 29ra30 (2010). 10.1126/scitranslmed.3000231

24. Florio, M., Gunasekaran, K., Stolina, M., Li, X., Liu, L., Tipton, B., Salimi-Moosavi, H., Asuncion, F. J., Li, C., Sun, B., Tan, H. L., Zhang, L., Han, C. Y., Case, R., Duguay, A. N., Grisanti, M., Stevens, J., Pretorius, J. K., Pacheco, E., Jones, H., Chen, Q., Soriano, B. D., Wen, J., Heron, B., Jacobsen, F. W., Brisan, E. & Ke, H. Z. A bispecific antibody targeting sclerostin and DKK-1 promotes bone mass accrual and fracture repair. Nat. Commun. 7, 11505 (2016). 10.1038/ncomms11505

25. Winkler, D. G., Sutherland, M. K., Geoghegan, J. C., Yu, C., Hayes, T., Skonier, J. E., Shpektor, D., Jonas, M., Kovacevich, B. R., Staehling-Hampton, K., Appleby, M., Brunkow, M. E. & Latham, J. A. Osteocyte control of bone formation via sclerostin, a novel BMP antagonist. EMBO J. 22, 6267–6276 (2003). 10.1093/emboj/cdg599

26. Duchamp de Lageneste, O., Julien, A., Abou-Khalil, R., Frangi, G., Carvalho, C., Cagnard, N., Cordier, C., Conway, S. J., & Colnot, C. (2018). Periosteum contains skeletal stem cells with high bone regenerative potential controlled by Periostin. Nature communications, 9(1), 773. 10.1038/s41467-018-03124-z

27. Ortinau, L. C., Wang, H., Lei, K., Deveza, L., Jeong, Y., Hara, Y., Grafe, I., Rosenfeld, S. B., Lee, D., Lee, B., Scadden, D. T., & Park, D. (2019). Identification of Functionally Distinct Mx1+αSMA+ Periosteal Skeletal Stem Cells. Cell stem cell, 25(6), 784–796.e5. 10.1016/j.stem.2019.11.003

28. Jeffery, E. C., Mann, T. L. A., Pool, J. A., Zhao, Z., & Morrison, S. J. (2022). Bone marrow and periosteal skeletal stem/progenitor cells make distinct contributions to bone maintenance and repair. Cell stem cell, 29(11), 1547–1561.e6. 10.1016/j.stem.2022.10.002

29. Lin, T. Y., Gerber, T., Taniguchi-Sugiura, Y., Murawala, P., Hermann, S., Grosser, L., Shibata, E., Treutlein, B., & Tanaka, E. M. Fibroblast dedifferentiation as a determinant of successful regeneration. Dev cell, 56(10), 1541–1551.e6. (2021). 10.1016/j.devcel.2021.04.016

30. Schindeler, A., Morse, A., Peacock, L. et al. Rapid cell culture and pre-clinical screening of a transforming growth factor-β (TGF-β) inhibitor for orthopaedics. BMC Musculoskelet. Disord. 11, 105 (2010). 10.1186/1471-2474-11-105

31. Wu, M., Wu, S., Chen, W. et al. The roles and regulatory mechanisms of TGF-β and BMP signaling in bone and cartilage development, homeostasis and disease. Cell Res. 34, 101–123 (2024). 10.1038/s41422-023-00918-9

32. Wei, E., Hu, M., Wu, L. et al. TGF-β signaling regulates differentiation of MSCs in bone metabolism: disputes among viewpoints. Stem Cell Res. Ther. 15, 156 (2024). 10.1186/s13287-024-03761-w

33. Livak, K. J. & Schmittgen, T. D. Analysis of relative gene expression data using real-time quantitative PCR and the 2(-Delta Delta C(T)) method. Methods 25, 402–408 (2001). 10.1006/meth.2001.1262

34. Nowoshilow, S., Schloissnig, S., Fei, J. F., Dahl, A., Pang, A. W. C., Pippel, M., Winkler, S., Hastie, A. R., Young, G., Roscito, J. G., Falcon, F., Knapp, D., Powell, S., Cruz, A., Cao, H., Habermann, B., Hiller, M., Tanaka, E. M. & Myers, E. W. Author Correction: The axolotl genome and the evolution of key tissue formation regulators. Nature 559, E2 (2018). 10.1038/s41586-018-0141-z

35. Schloissnig, S., Kawaguchi, A., Nowoshilow, S., Falcon, F., Otsuki, L., Tardivo, P., Timoshevskaya, N., Keinath, M. C., Smith, J. J., Voss, S. R. & Tanaka, E. M. The giant axolotl genome uncovers the evolution, scaling, and transcriptional control of complex gene loci. Proc. Natl Acad. Sci. USA 118, e2017176118 (2021). 10.1073/pnas.2017176118

36. Dobin, A., Davis, C. A., Schlesinger, F., Drenkow, J., Zaleski, C., Jha, S., Batut, P., Chaisson, M. & Gingeras, T. R. STAR: ultrafast universal RNA-seq aligner. Bioinformatics 29, 15–21 (2013). 10.1093/bioinformatics/bts635

37. Hao, Y., Hao, S., Andersen-Nissen, E., Mauck, W. M. 3rd, Zheng, S., Butler, A., Lee, M. J., Wilk, A. J., Darby, C., Zager, M., Hoffman, P., Stoeckius, M., Papalexi, E., Mimitou, E. P., Jain, J., Srivastava, A., Stuart, T., Fleming, L. M., Yeung, B., Rogers, A. J., McElrath, J. M., Blish, C. A., Gottardo, R., Smibert, P. & Satija, R. Integrated analysis of multimodal single-cell data. Cell 184, 3573–3587.e29 (2021). 10.1016/j.cell.2021.04.048

38. Korsunsky, I., Millard, N., Fan, J., Slowikowski, K., Zhang, F., Wei, K., Baglaenko, Y., Brenner, M., Loh, P. R. & Raychaudhuri, S. Fast, sensitive and accurate integration of single-cell data with Harmony. Nat. Methods 16, 1289–1296 (2019). 10.1038/s41592-019-0619-0

39. Angerer, P., Haghverdi, L., Büttner, M., Theis, F. J., Marr, C. & Buettner, F. destiny: diffusion maps for large-scale single-cell data in R. Bioinformatics 32, 1241–1243 (2016). 10.1093/bioinformatics/btv715

40. Schreiber, T. Measuring information transfer. Phys. Rev. Lett. 85, 461–464 (2000). 10.1103/PhysRevLett.85.461

41. Shen, W. K., Chen, S. Y., Gan, Z. Q., Zhang, Y. Z., Yue, T., Chen, M. M., Xue, Y., Hu, H. & Guo, A. Y. AnimalTFDB 4.0: a comprehensive animal transcription factor database updated with variation and expression annotations. Nucleic Acids Res. 51, D39–D45 (2023). 10.1093/nar/gkac907

42. Kanehisa, M. & Goto, S. KEGG: kyoto encyclopedia of genes and genomes. Nucleic Acids Res. 28, 27–30 (2000). 10.1093/nar/28.1.27

43. Kanehisa, M., Sato, Y., Furumichi, M., Morishima, K. & Tanabe, M. New approach for understanding genome variations in KEGG. Nucleic Acids Res. 47, D590–D595 (2019). 10.1093/nar/gky962

44. Korotkevich, G., Sukhov, V., Budin, N., Shpak, B., Artyomov, M. N. & Sergushichev, A. Fast gene set enrichment analysis. bioRxiv 060012 (2021). 10.1101/060012

45. Blakney, A. K., McKay, P. F., Hu, K., Samnuan, K., Jain, N., Brown, A., Thomas, A., Rogers, P., Polra, K., Sallah, H., Yeow, J., Zhu, Y., Stevens, M. M., Geall, A. & Shattock, R. J. Polymeric and lipid nanoparticles for delivery of self-amplifying RNA vaccines. J. Control Release 338, 201–210 (2021). 10.1016/j.jconrel.2021.08.029

46. Hutchison, C., Pilote, M. & Roy, S. The axolotl limb: a model for bone development, regeneration and fracture healing. Bone 40, 45–56 (2007). 10.1016/j.bone.2006.07.022

47. Berg, S., Kutra, D., Kroeger, T., Straehle, C. N., Kausler, B. X., Haubold, C., Schiegg, M., Ales, J., Beier, T., Rudy, M., Eren, K., Cervantes, J. I., Xu, B., Beuttenmueller, F., Wolny, A., Zhang, C., Koethe, U., Hamprecht, F. A. & Kreshuk, A. ilastik: interactive machine learning for (bio)image analysis. Nat. Methods 16, 1226–1232 (2019). 10.1038/s41592-019-0582-9

